# Molecular signatures of altered energy metabolism and circadian rhythm perturbations in a model of extra-nigral Synucleinopathy

**DOI:** 10.1101/2024.09.27.615378

**Authors:** Lin Lin, Nanna M. Jensen, Sara A. Ferreira, Alberto Delaidelli, Diana Gomes Moreira, Ole A. Ahlgreen, Mette Richner, Poul H. Sorensen, Marina Romero-Ramos, Christian B. Vægter, Poul H. Jensen, Ian R. Mackenzie, Jens R. Nyengaard, Asad Jan

**Affiliations:** Department of Biomedicine, Aarhus University, Høegh-Guldbergs Gade 10, DK-8000 Aarhus Denmark; Danish Research Institute of Translational Neuroscience (DANDRITE), Aarhus University, Ole Worms Allé 3, DK-8000 Aarhus C, Denmark; Department of Clinical Medicine, Aarhus University, Palle Juul-Jensens Boulevard 35, DK-8200 Aarhus, Denmark; Core Center for Molecular Morphology, Section for Stereology and Microscopy Department of Clinical Medicine, Aarhus University, DK-8200 Aarhus N, Denmark; Department of Pathology and Laboratory Medicine, 2211 Wesbrook Mall, Vancouver, BC V6T 2B5, Canada; British Columbia Cancer Research Centre, Vancouver, BC V5Z 1L3 Canada

**Keywords:** Parkinson Disease, Αlpha-Synuclein, Spatial Transcriptomics, Mitochondrial Metabolism, Circadian Rhythm

## Abstract

A pathological role of alpha-Synuclein (aSyn) aggregation in the central nervous system (CNS) is a recognized feature in Parkinson disease (PD) and related neurodegenerative conditions termed synucleinopathies. In order to characterize the cellular response in CNS to incipient and advanced aSyn pathology, we applied spatial transcriptomics on brain sections derived from a transgenic mouse model (M83^+/+^ line, *Prnp-SNCA*A53T*) in which aSyn aggregation was induced in a prion-like fashion through hindlimb intramuscular delivery of pre-formed fibrillar (PFF) murine aSyn. Our spatially-resolved transcriptomics (ST) data point to unique perturbations in brain energy metabolism during the progression of aSyn pathology, such that the early stage of aSyn aggregate pathology activates molecular pathways controlling metabolic flux through glycolysis, oxidative phosphorylation and fatty acid metabolism. In contrast, the ST data indicate a profound decline in mitochondrial metabolism in the brains of symptomatic animals with advanced aSyn pathology. The latter stage was also associated with drastic reduction in mRNA translation machinery, along with aberrant expression of molecular drivers involved in RNA splicing and inflammatory response. Intriguingly, our ST data also point to perturbed regulation of circadian rhythm, was corroborated by increased immunodetection of CREB-binding protein (a modulator of core clock machinery) in the brains of symptomatic animals, and transcriptional upregulation of *CREBBP* in 4 independent PD microarray datasets. Collectively, we anticipate that our findings offer novel opportunities in knowledge translation for mechanism-based drug discovery and biomarkers in neurodegenerative synucleinopathies.

## INTRODUCTION

Cellular alpha-Synuclein (aSyn; gene symbol *SNCA*) pathology in the nervous system is a hallmark feature of idiopathic Parkinson disease (PD) and related synucleinopathies ^1,2^. Under physiological conditions, aSyn protein has been implicated in facilitating the assembly of synaptic components mediating vesicle release and as a chaperone ^3,4^. However, in pathological states, aSyn is prone to self-assemble in a prion-like fashion and accumulates in neuronal perikarya and neuropil in the form of Lewy-related aSyn pathology ^5^. In the context of neurodegeneration in PD, studies suggest that the process of aSyn aggregation adversely affects neuronal survival by triggering mitochondrial impairment, endoplasmic reticulum (ER) stress, defects in protein sorting and autophagy, perturbed redox homeostasis and neuroinflammation ^1,4^. This notion is also supported by the genetic association of rare *SNCA* variants (A30P, E46K, or A53T) and gene duplications/triplications to familial forms of early-onset aggressive parkinsonism ^1,2^. Moreover, transgenic and/or ectopic overexpression of aSyn using viral vectors is a widely used method in model development for PD and related diseases, both for investigating the disease mechanisms as well as for evaluating the disease modifying effects of candidate therapeutic molecules ^6,7^.

The progressive degeneration of dopaminergic neurons in the midbrain substantia nigra, *pars compacta* (SN*pc*) is considered to be the neural substrate for clinical parkinsonism, exemplified by PD, in the vast majority of cases ^1,2^. However, there is growing consensus that the nigro-centric view is not sufficient to account for the heterogeneity in clinical presentation of PD, especially in the context of prodromal non-motor symptomatology affecting the sense of smell, sleep and autonomic function ^5,8^. In this regard, seminal neuropathological studies by Braak and colleagues, and corroborated by others, suggest that pathological aSyn deposition in extra-nigral brain regions appears long before the affection of SN*pc* ^9–11^. In particular, the neuronal populations within the brainstem reticular formation (BRF, in pons and medulla), dorsal motor nucleus of vagus (dmX) and locus coeruleus (LC) bear the brunt of cellular aSyn pathology in prodromal PD (Braak stages I-II)-^9–11^. These findings have ushered novel efforts in preclinical model development for prodromal PD symptomatology, for instance aSyn aggregation in dmX for studying the gut-brain axis and autonomic dysfunction ^12–14^.

In a mouse model of synucleinopathy (M83 line- *Prnp-SNCA*A53T*, overexpressing the aggregation prone human mutant A53T) ^15^, we and other have shown that exogenous delivery of aggregated (PFF) aSyn using peripheral (intramuscular) route reproducibly induces *de novo* aSyn aggregation in the nervous system ^16–20^. These studies show that the spatiotemporal propagation of PFF-induced aSyn pathology in the brains of these rodents is not random, such that the paramedian nuclei in brainstem (GRN, vestibular nuclei and mesencephalic PAG) are among the earliest neuronal populations which are preferentially affected during the prodromal (non-symptomatic stage) ^17,18,21^. Notably, these extra-nigral regions, especially GRN and PAG, have also been reported to harbor substantial pathological aSyn accumulation in PD, with potential implications for non-motor symptomatology ^9,10,22^. In addition, this rodent model exhibits aSyn accumulation in the white matter (oligodendrocytes and myelin tracts), a feature reminiscent of Lewy-related aSyn pathology seen in PD and other synucleinopathies ^5,18,22^. It is worth mentioning that the pathological involvement of substantia nigra (SN), hypothalamus and parts of thalamus is rarely observed in this rodent model ^15,18–21^. Nevertheless, this experimental paradigm has been informative in studying the consequences of extra-nigral aSyn pathology in brainstem. For instance, long before the onset of movement disability, this model exhibits alterations in the patterns of brain activity ^23^, defects in the nerve conduction ^17^, pain-related behaviors ^17^, and aberrations in the neuronal expression of tissue homeostatic factors, which in some instances are corroborated by histological studies in post-mortem PD brains ^24,25^.

Hence, we hypothesized that the distinct phases of aSyn aggregation (ie. prodromal and symptomatic) in the brains of these rodents offer a window of opportunity for studying the cellular effects of incipient and advanced aSyn pathology in the nervous system. In other words, we postulated that: i) the prodromal aSyn pathology in the nuclei of BRF (GRN and PAG) ^18^ will allow us to decipher the initial cellular response in these extra-nigral neuronal populations to *de novo* aSyn aggregation, and ii) the progression phase (associated with the symptomatic stage) ^18–20^ will potentially yield valuable insights into the mechanism(s) dictating cell-to-cell transmission and neuronal and/or glial dysfunction. Thus, we harnessed the utility of spatial transcriptomics (ST) to obtain spatially resolved brain gene expression profiles associated with incipient and advanced stages of cellular aSyn pathology in this rodent model. Differential gene expression (DGE) profiling, including the analyses for transcription factors (TFs) that control the sets of DGE profiles, and pathway enrichment analyses revealed that the prodromal phase is associated with the upregulation of transcripts controlling mitochondrial metabolism in the disease-affected regions. In contrast, during the symptomatic phase, the ST data suggest profound decline in brain energy metabolism, reflected by the downregulation of transcripts encoding factors involved in glycloysis, oxidative phosphorylation and fatty acid metabolism. This latter phase was also associated with altered expression of transcripts belonging to ribosomal mRNA translation machinery, and pathways of neuroinflammation and circadian rhythms. We also present histological analyses assessing the expression of CREB-binding protein (CREBBP/CBP) in brain sections from the rodent model and post-mortem PD midbrain, which was identified as a differentially expressed transcript in the ST data and in PD microarray studies. Collectively, these data point to unique molecular signatures of cellular dysfunction induced by progressive aSyn aggregate pathology in the nervous system, and will serve as a valuable resource for translational research in PD and related diseases.

## MATERIALS AND METHODS

### Generation and characterization of mouse aSyn fibrils

Mouse aSyn fibrils (PFF) were prepared and characterized *in vitro*, essentially as described ^17,18^. Briefly, full length recombinant (wild type) mouse αSyn was expressed in BL21(DE3) competent cells and purified using reverse phase chromatography. Then, purified recombinant monomeric (ie., non-aggregated) α-Syn (10 mg/ml) was incubated at 37°C in phosphate-buffered saline (PBS, pH 7.4) with continuous shaking at 1050 r.p.m. in a tabletop microtubes shaker (Eppendorf). The PFF were collected by centrifugation (15,600g at 25°C for 30 min), and then re-suspended in PBS. Protein concentration was determined by the BCA assay (Pierce) and a stock solution consisting of 2 mg/mL protein was prepared (in PBS). Subsequently, PFF were sonicated for 20 minutes using a Branson 250 Sonifier at 30% intensity, and then aliquoted and frozen at□−□80 °C until further use. The purity of these fibrillar preparations, their biophysical characterization and biological activity is described elsewhere ^17,18^.

### Animal Studies

#### Hindlimb intramuscular PFF aSyn injection

Transgenic M83 mice [B6;C3-Tg(*Prnp- SNCA*A53T*)83Vle/J]-^15^ were housed at the skou animal facility at Aarhus University in accordance with Danish regulations and the European Communities Council Directive for laboratory animals (license # 2017-15-0201-01203 issued to PHJ, co-author). The animals were housed under a 12 hours light/dark cycle and fed with regular chow diet *ad libitum*. Adult homozygous M83^+/+^ mice (12-14 weeks of age) were bilaterally inoculated with a single injection (5 μl) of recombinant mouse aSyn PFF (1 μg/μl; n=6) into the hindlimb biceps femoris, using a 10-μL Hamilton syringe with a 25-gauge needle as described in the foundational study ^18^. Age-matched controls were injected with PBS (5 µl, bilaterally; n=2). The experiments included both male and female mice.

#### Collection of brains and preparing tissue sections

Mice were transcardially perfused with ice-cold PBS supplemented with 10 U/ml Heparin (Sigma, #H3149). Brains were removed from the skull and immediately transferred on ice-cold PBS. Then, the brains hemisphere were embedded in Tissue-Tek OCT (Sakura, #4583), immediately frozen by liquid nitrogen bath and kept at −80°C until cryosectioning. For ST and related analyses in the present study, the OCT embedded was hemisphere was initially trimmed from the midline, covering ∼50 µm, to expose the paramedian nuclei (GRN and PAG; see note on Mouse Neuroanatomical Topography below) ^26^. Then, five 10 µm thick sagittal serial sections were obtained, with the cryostat chamber maintained at −20°C. The outer four sections, placed on the Superfrost Plus slides (VWR, #631-0108), were stored −80°C and reserved for the immunofluorescence (IF) analyses. The central section, used for ST, was placed within the capture area (6.5 x 6.5 mm) on the Visium Spatial Gene Expression slides (10x Genomics, #PN-1000185), covering brain regions of interest (paramedian pons and midbrain, cerebellum, thalamus, hypothalamus and cortex) ^18,20,21^.

### Spatial Transcriptomics

#### Spatial library preparation

Spatial bar coded ST library from fresh frozen brain sections was prepared using Visium Spatial Gene Expression Slide & Reagent Kit (10x Genomics, #PN-1000184), according to the manufacturer’s provided instructions (CG000240 Rev C). The sections were fixed in pre-chilled (−20°C) methanol (Sigma, #34860) for 10 min at −20°C. Sections were counterstained with hematoxylin (Vector Labs, #H-3401) for 5 min at room temperature (RT) and whole slide digital scans were obtained using the Olympus VS120 upright microscope in the brightfield mode (20X magnification). Samples were permeabilized for 18 min, based on the optimal time determined in the tissue optimization procedure (10x Genomics, #1000193 & Cat#CG000238 Rev D). Spatial cDNA libraries were prepared according to the manufacturer’s guidelines for the procedure (CG000240 Rev C: reverse transcription, second strand synthesis, cDNA amplification and adapter ligation). The resulting libraries were pooled and sequenced on an MGI G400 sequencer, followed by de-multiplexing using the library index sequences.

#### Sequencing Data Alignment and Normalization

Sequencing data in FASTQ format were aligned to the reference genome (refdata-gex-mm10-2020-A) using Space Ranger software (10x Genomics, version 1.3.0). Briefly, spots under tissues were selected in LoupeBrowser, json files were then exported and used for alignment with Space Ranger. The Seurat package was utilized for data normalization and annotation of each cryosection. Raw count inputs were normalized using the SCTransform normalization method. Following normalization, principal component analysis (PCA) was performed for linear dimension reduction, selecting the first 30 principal components (PCs) to create a shared nearest neighbor (SNN) network. The same PCs were employed for visualization using Uniform Manifold Approximation and Projection (UMAP) and t-Distributed Stochastic Neighbor Embedding (t-SNE) algorithms.

#### Spatial annotation

Clusters within each cryosection were annotated by calculating cluster-specific markers using the “FindAllMarkers” function. This determined log fold change, percentage of expression within and outside the target cluster, and p-values for the Wilcoxon rank-sum test. A log2 fold-change (FC) threshold of ≥±0.25 and an adjusted p-value ≤0.05 were considered significant. By comparing identified marker genes with canonical region marker genes, each cluster was assigned an ST regional annotation.

#### Merging data and unsupervised Clustering

Individual Seurat objects for each cryosection were combined into a single, integrated dataset. Unsupervised clustering was conducted, involving PCA to identify major axes of variation within the data. The first 20 PCs were selected for further analysis, generating an SNN network to reveal the data’s underlying structure. These 20 PCs were used for visualization, applying dimensionality reduction techniques such as UMAP and t-SNE to represent and interpret high-dimensional data more comprehensibly.

#### Correlation analyses

To assess relationships between different regions within each region/cryosection, the average gene expression for all genes in each region was calculated. Pair-wise correlation coefficients between all pairs of regions were computed based on their average gene expression profiles using the ‘cor’ function from the ‘stats’ package. A Euclidean distance-based dissimilarity matrix was constructed from the resulting correlation matrix. Hierarchical clustering was then performed on the dissimilarity matrix using the complete linkage method. This approach identified patterns and relationships between distinct regions across cryosections based on their gene expression similarities, providing insights into the underlying biological processes and molecular signatures.

#### Pair-wise comparison of Differential Gene Expression

Differentially expressed genes between two groups were identified using the ‘FindMarkers’ function in the Seurat package. This function calculates the log2 fold change (FC) in gene expression, the percentage of expression within and outside the target cluster, and p-values using the likelihood-ratio test. DEGs were considered significant if they met the following criteria: a log2 FC threshold of ≥±0.25 and an adjusted p-value of ≤0.05.

#### Gene Set Enrichemnt (GSE) Analyses

To investigate the functional relevance of gene expression patterns in our dataset, we employed two distinct strategies for gene set analysis. The gene sets were collected from the Molecular Signatures Database (msigdbr version 7.5.1; http://bioinf.wehi.edu.au/software/MSigDB/) using the msigdbr package, enabling us to identify enriched pathways and cellular processes across different regions and time points.

The first strategy involved performing Gene Set Variation Analysis (GSVA) on the average expression for each cluster. The average expression was calculated for each region in different cryosections. GSVA was conducted using the GSVA R-package (version 1.46.0), which converts the gene-by-cluster matrix into a gene-set-by-cluster matrix, providing a pathway-centric perspective on the data.

The second strategy employed the ‘AddModuleScore’ function from the Seurat package to calculate module scores for individual gene sets. This approach enabled the computation of module scores for each spot in the dataset, offering a more granular view of gene set activity across the spatial transcriptomic landscape. Together, these complementary strategies allowed for a comprehensive understanding of the biological processes and molecular pathways underlying the observed gene expression patterns in the ST dataset.

#### Data Visualization

A range of visualization techniques were employed to effectively represent the ST data. We generated UMAP and spatial plots, violin plots, and gene expression dot plots using the ‘DimPlot’, ‘SpatialDimPlot’, ‘SpatialPlot’, ‘VlnPlot’, and ‘DotPlot’ functions within the Seurat package. Additionally, heatmaps were created utilizing the ‘heatmap’ function in the stats package (version 4.0.3). Genes responded in different regions at different time points were analyzed and visualized with UpsetR package. ST data were also integrated and visualized as KEGG pathway maps using Pathview R package ^27^.

### Histological Analyses

#### Immunofluorescence (IF) analyses of mouse brain sections

IF was performed on 10 µm thick fresh frozen sections, after drying (5 min, at 37°C in a non-humid oven) and gentle fixation with ice-cold 4% Paraformaldehyde (Electron Microscopy Sciences, #15700; 15 min, at 4°C). Nonspecific binding was blocked by incubating the sections in 5% normal donkey serum in Tris-buffered saline (1 hour, RT). Then, the sections were incubated (overnight, at 4°C) with the following primary antibodies (in PBS containing 0.3% Triton-X and 0.5% bovine serum albumin-BSA): phospho-S129 aSyn (rabbit mAb EP1536Y, Abcam, #ab51253-dilution: 1:1000), Creb-binding protein (CREBBP/CBP; rabbit mAb EPR23418-23 Abcam, #ab253202-dilution: 1:500) and Rho Associated Coiled-Coil Containing Protein Kinase 2 (ROCK2, rabbit pAb, LSBio#LS-C3334228-dilution: 1:250). IF detection was performed by Alexa-Fluor488 fluorophore conjugated secondary antibody (Thermo Fisher #A-11008-dilution: 1:1000).

#### Mouse Neuroanatomical Topography

Panoramic images from the digital whole slide scans were mapped onto Mouse Brain Atlas (Paxinos and Franklin’s The Mouse Brain in Stereotaxic Coordinates, 4^th^ Edition)-^26^. Information about neuroanatomical tracts and nuclei was primarily derived from The Mouse Nervous System (1^st^ Edition)-^28^.

#### Human tissue processing and Immunohistochemistry (IHC) analyses

IHC data from post-mortem control and PD midbrain sections were acquired by using p-aSyn (p-S129) antibody (rabbit mAb EP1536Y Abcam, #ab51253-1:1000) and anti-CREBBP (rabbit mAb EPR23418-23 Abcam, #ab253202-1:500) essentially as described ^25^. Five micrometer formalin-fixed paraffin embedded sections were kindly provided by IRM (co-author), after the study approval by the University of British Columbia Ethics committee (Supplementary Table S1). IHC on brain sections was performed after deparaffinization and antigen retrieval. For destaining/bleaching of neuromelanin in the substantia nigra (in p-aSyn S129 IHC studies), a slightly modified IHC protocol was used ^18,25^. Briefly, slides with mounted sections were incubated in a 60°C degrees oven for 30 minutes and then were transferred into ambient distilled water. Then, the slides were placed in 0.25% potassium permanganate solution for 5 minutes. Subsequently, the slides were rinsed with distilled water. This was followed by incubation in 5% oxalic acid until section became clear. A final rinse in distilled water was performed before proceeding with the routine IHC staining on serial sections as described below. Tissue sections were incubated with the primary antibodies and immunodetection was performed using the alkaline phosphatise conjugated streptavidin-biotin ABC kit (Vector Labs, #AK-5000). Sections were counterstained with hematoxylin (Vector Labs, #H-3401).

#### Image analyses- Mouse brain sections

High resolution IF views of the mouse brain sections were obtained using Olympus VS120 digital slide scanner equipped for fluorescence single-band emitters for Hoechst, FITC, Cy3 and Cy5 (AU, Denmark). Slide scans were imported into Qupath (v. 0.5.1) ^29^ and entire tissue regions were outlined based on thresholding for the nuclear DAPI staining. Regions of interest (ROI) consisting of GRN (pons), PAG (midbrain), deep cerebellar nuclei (DCN) and mediodorsal (MD) thalamic nuclei were manually outlined in each section, using the Mouse Brain Atlas (Paxinos and Franklin’s The Mouse Brain in Stereotaxic Coordinates, 4^th^ Edition)-^26^ as a guide. ROI were segmented using the Qupath cell detection plugin on the DAPI channel. Phospho-aSyn (S129) threshold was computed as 10 x median IF intensity in the entire tissue region. This threshold was applied in Pixel Classification thresholder on the detection channel, using full image resolution and no prefiltering/smoothing. The thresholder was then used to compute phospho-aSyn (S129)-positive area as percentage (%) of the total ROI area.

For analyzing the IF staining for CREBBP, automated thresholding was performed on the detection channel (script for graphical user interface available at https://github.com/iviecomarti/GUI_AutoTH_QuPath/blob/main/gui/gui_AutoTH_QuPath.groovy). Thresholding was performed on the entire tissue region using the “Moments” method to get a threshold value as output. Subsequently, this threshold was multiplied by 1.2 and put into a Single Measurements Classifier on the CREBBP channel to compute CREBBP-positive cells, defined as having a mean intensity in the nucleus above the set threshold. Results were plotted as CREBBP-positive cells in %of total cells in the ROI. For the analysis of ROCK2 IF staining, median ROCK2 intensity was computed in each ROI and multiplied by 1.5 to get the threshold. This threshold was put into a Single Measurements Classifier on the detections channel to compute ROKC2-positive cells, defined as having a mean intensity in the cytoplasm above the set threshold. Results were graphed as ROCK2-positive cells in % of total cells in the ROI.

#### Image analyses- Human brain sections

Anonymized whole slide digital panoramic images of the tissue sections for IHC analyses were acquired using a Leica Aperio slide scanner (UBC, Canada). Slide scans from PD cases and non-neurodegenerative controls were imported into Qupath (v. 0.5.1) ^29^, and ROIs of approx. 720 x 500 µm were defined for the SN and PAG in both hemispheres (one in each hemisphere). All analysis was performed blinded to the diagnosis of the cases and CREBBP staining was analyzed before assessing the Phospho-aSyn (S129) stained sections. For the analyses of CREBBP IHC staining, large blood vessels and neuromelanin was first excluded from the ROIs. Then, stain vectors were estimated using the auto function in Qupath. Positive cell detection was run on the ROIs with nuclei detected by hematoxylin stain and positive cells defined by a mean optical density of nuclear DAB staining above 0.2. Results were graphed as CREBBP-positive cells in % of the total cells each ROI. For the phospho-aSyn (S129) IHC analyses, LBs were manually counted in each ROI and normalized to the total ROI area to obtain the number of LBs per mm2 of the tissue ^24,25^.

#### Statistics

The data were statistically analyzed in Graphpad Prism software version 8, and the final graphs were prepared in Graphpad or Microsoft Excel. Statistical significance in datasets was calculated and indicated in the relevant figure legends. P-values were set at: *p<0.05, **p<0.01, ***p<0.001, ****p<0.0001.

#### Data and code availability

All of the data generated and analyzed during this study are included in the main manuscript and the associated supplementary files. The processed and raw data generated during this study have been made publicly accessible through the GEO repository (accession number: GSE274605). For the visualization and interactive exploration of the ST data, we have created a dedicated online interface (URL: https://dreamapp.biomed.au.dk/PD_spatial_mouse_DB, and a short User Guide is provided in the Supplementary Information. This interactive platform has been developed and customized using the ShinyCell package, allowing the users to engage with the data more effectively.

## RESULTS

### Intramuscular PFF aSyn delivery induced progressive aSyn aggregation in the GRN and PAG of M83^+/+^ mice

In this study, we wanted to decipher the molecular signatures associated with the progression of cellular aSyn pathology in the nervous system, which is a crucial factor implicated in the pathogenesis of neurodegeneration in PD and related synucleinopathies ^1,2^. Specifically, we applied ST to sagittal brain sections obtained from the M83^+/+^ mice, following intramuscular delivery of PFF aSyn at: i) an early stage (ES, days post-injection: DPI-45), while the mice were freely moving with lack of gross movement disability, albeit a mild degree of hindlimb clasping ^18^ and ii) a late stage (LS, DPI-75), when the mice started to exhibit the onset of a rapidly progressive movement disability. We chose to analyze the sagittal brain sections for the purpose of ST (in contrast to the region-limited coronal sections, as used in the previous studies ^17,18,21^), since: i) it allowed us to capture the brain regions of interest affected in the prodromal phase, especially the paramedian nuclei of reticular formation (GRN and PAG) ^17,18,21^, ii) the large ascending and descending tracts of the white matter, affected by aSyn aggregation in this model ^18^, are represented throughout the entire length of the section, and iii) a broader caudo-rostral “snapshot” of the whole brain can be obtained in one section, which is not possible in a coronal section. Prior to processing the sections for ST, we assessed the extent of aSyn aggregate pathology by immunodetection of aSyn phosphorylation on serine residue S129 (p-aSyn, S129), which is one of the widely used biochemical marker in PD neuropathology studies ^30–32^. In line with the previous observations (based on coronal sections) ^17,18,21^, we observed a progressive accumulation of aggregated aSyn in the GRN (pons) and PAG (midbrain) of PFF aSyn-injected M83^+/+^ mice, and to a lesser extent in the DCN (cerebellum) and MD thalamic nuclei (Fig. 1A-C).

**Figure 1.**
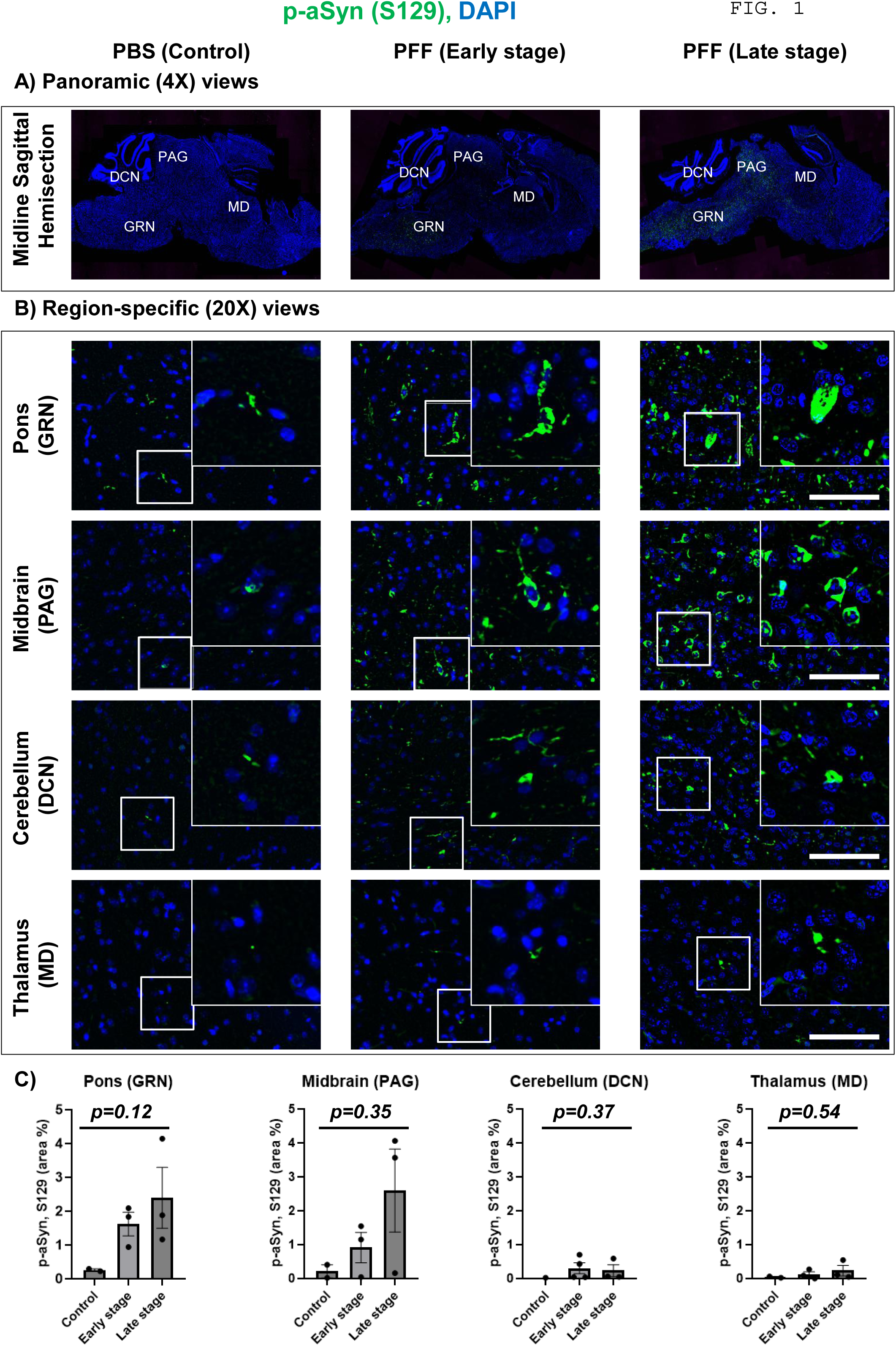
Immunofluorescence (IF) detection of phosphorylated alpha-synuclein (p-aSyn, S129) in brains of M83^+/+^ mice. **(A)** Representative low magnification (4X) panoramic images of sagittal brain sections from controls (PBS, Days post-injection, DPI-75) and PFF aSyn-injected M83^+/+^ mice (Early stage, DPI-45; and Late stage, DPI-75). The labels in CAPITAL LETTERS indicate specific brain regions examined for the IF analyses (in B). **(B)** Representative (20X) images showing p-aSyn (S129) IF in the gigantocellular nuclei (GRN, in pons), periaqueductal grey (PAG, in midbrain), deep cerebellar nuclei (DCN, in cerebellum) and mediodorsal nuclei (MD, in thalamus). The inset show 63X magnified views from the regions. Scale bar= 50 µm. **(C)** Bar graphs depicting quantification of p-aSyn (S129) IF in GRN, PAG, DCN and DM of the experimental cohorts, as indicated. Error bars depict Mean IF intensity ± SD as % of total area (PBS, n=2; DPI-45, n=3 and DPI-75, n=3; See Methods). Statistics in Fig. 1C: Kruskal-Wallis ANOVA (p-values on graphs), Dunn’s multiple comparisons (non-significant).

### The clustering of ST profiles corresponded to the stage of aSyn pathology and distinct neuroanatomical annotations in mouse brain

Next, we employed a pipeline for annotating the ST data onto the histological map of the sagittal brain sections (see Methods) obtained from controls (PBS, n=2) and PFF aSyn-injected M83^+/+^ mice (ES, n=3; LS, n=3). We identified 16833 unique protein coding transcripts in the ST data, which clustered distinctively between the experimental groups (Fig. 2A, UMAP plots), and in anatomical subregions (Fig. 2B,UMAP plots). The latter analyses incorporate the spatial elements reflecting gross anatomical identity of the grey matter regions, with guidance from Mouse Brain Atlas (namely: pons, midbrain, cerebellum, thalamus, hypothalamus and cortex including hippocampal formation-Fig. 2C-D) ^26,28^, and confirmed by specific sets of regionally-enriched transcripts in the mouse brain (https://www.proteinatlas.org/humanproteome/brain/mouse+brain) ^33^. To avoid ambiguity between the grey vs. non-grey matter, the ST annotations were also adapted to account for large white matter tracts and cerebral ventricular system (choroid plexus), as illustrated below.

**Figure 2.**
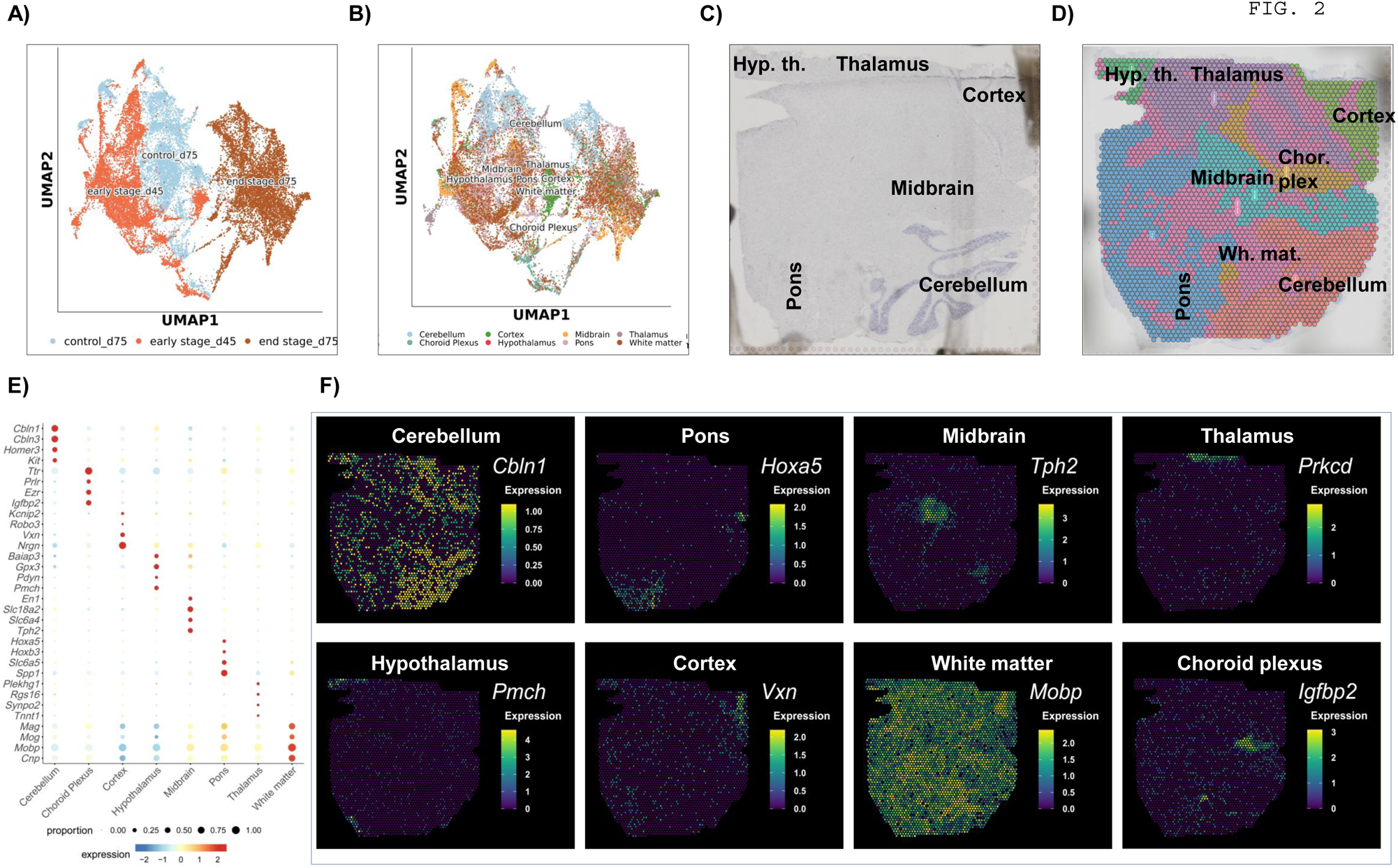
Overview of ST annotations using Neuroanatomical topography and regionally enriched transcripts in cohorts of M83 M83^+/+^ mice. **(A-B)** UMAP plots depicting distinct clustering of ST profiles in the brains of experimental M83^+/+^ cohorts in relation to the stage of aSyn pathology (in A) and 8 ST annotations (in B). **(C-D)** Representative low magnification (4X) panoramic images of a sagittal brain section showing grey matter regions (in C, hematoxylin counterstain) and spatial map (in D, with the addition of white matter and choroid plexus annotations guided by the abundance of regional markers in E; also see Methods). **(E-F)** Illustrations of regionally enriched markers used in delineating 8 ST annotations, with bubble plot (in E) and spatial map of select markers in a sagittal brain section. Spatial plots in D: *Cbln1* (cerebellum)*, Hoxa5* (Pons)*, Tph2* (midbrain)*, Prkcd* (thalamus)*, Pmch* (hypothalamus)*, Vxn* (cerebral cortex)*, Mobp* (white matter) and *Igfbp2* (choroid plexus). Notice the imperfect segregation of whiter matter in brainstem (pons and midbrain) compared to cortex.

In brief, the 8 ST annotation profiles exhibited regional enrichment of unique transcripts reflecting synaptic markers in grey matter (eg., *Homer3* in cerebellum, *Tph2* in midbrain and *Slc6a5* in pons), structural components in the white matter (eg., *Mobp*) and choroid plexus in the cerebral ventricles (eg., *Ttr*)-Fig. 2E. This was also reflected in the spatial plots (Fig. 2F), showing region-enhanced expression of *Cbln1* (cerebellum)*, Hoxa5* (pons)*, Tph2* (midbrain)*, Prkcd* (thalamus)*, Pmch* (hypothalamus)*, Vxn* (cerebral cortex)*, Mobp* (white matter) and *Igfbp2* (choroid plexus) ^33^. Admittedly, there were limitations to conclusively differentiate every spatial element (capture spot) into the grey and non-grey matter categories at the microscopic detail, especially in pons and midbrain (Fig. 2E). This is mainly due to two reasons: i) the applied technology (10x Genomics Visium v1) lacks the single-cell resolution, such that each spot in the capture area is 55 µm in diameter with an inter-spot center-to-center distance of 100 µm, and ii) the two regions (pons and midbrain) contain major cranial nerve nuclei and neuronal populations of the reticular formation which are embedded by dense clusters of white matter and traversed by large sensorimotor tracts ^28^. In contrast, the neuroanatomical regions where white matter elements (oligodendrocytes and myelin tracts) are predominantly organized in distinct tracts/bundles (cerebellum, cortex), or are completely lacking (choroid plexus), a clear grey vs. non-grey matter distinction was possible (Fig. 2E; compare cortex, with pons and mibrain). Thus, our ST annotation for the 8 regional profiles (Fig. 2B-D) reflects gross anatomical organization of brain (and not single-cell, microscopic organization), which is further fine-tuned by spatial expression pattern of the region-enriched transcripts (Fig. 2E-F) ^33^. With this limitation in perspective, below we describe the salient findings from the ST study in this rodent model of progressive aSyn pathology in the nervous system.

### ST unveiled unique molecular signatures associated with the early and late stages of aSyn pathology in the brains of M83^+/+^ mice

Next, we performed gene set enrichment (GSE) analyses for comparing the expression pattern of the transcripts belonging to specific cellular pathways, using HALLMARK and Kyoto Encyclopedia of Genes and Genomes (KEGG) gene sets, available on the MSigDB portal (https://bioinf.wehi.edu.au/software/MSigDB/). These analyses revealed that the 8 ST annotation profiles in general, and the disease affected profiles (pons, midbrain and white matter) in particular, exhibited a biphasic pattern in the expression of transcripts belonging to the pathways controlling energy metabolism. Specifically, the data suggest significant upregulation in the pathways controlling oxidative phosphorylation, glycolysis, fatty acid metabolism, cholesterol homeostasis, and mTORc1 signaling in association with ES aSyn pathology, followed by a drastic downregulation in these pathways by the LS (Fig. 3A). Moreover, throughout the grey matter annotations, significant transcriptional upregulation in unfolded protein response (a crucial biological process involved in proper intracellular sorting and/or degradation of misfolded protein, including aSyn) ^4^ was found. Intriguingly, the ES was also associated with augmentation in pathways controlling reactive oxygen species (ROS) metabolism and DNA repair, detected across the grey and white matter annotations (Fig. 3A).

**Figure 3.**
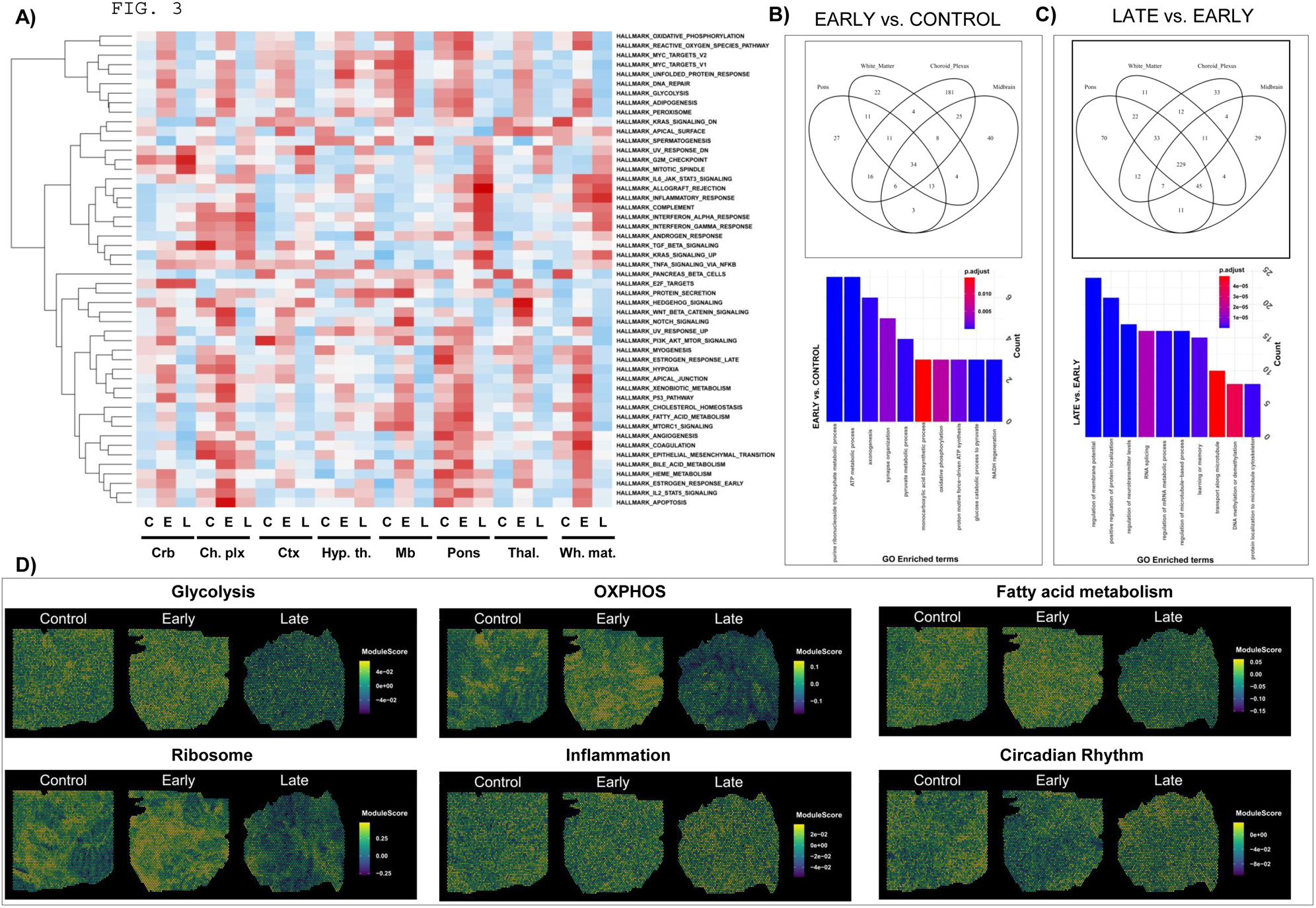
Overview of the HALLMARK gene sets and differential gene expression across the ST annotations in cohorts of M83^+/+^ mice. **(A)** Heatmap depicting the relative expression of HALLMARKS gene sets in pair-wise comparisons involving Early stage (E) vs. Controls (C) and Late stage (L) vs. Early stage (E) experimental cohorts of M83^+/+^ mice. Abbreviations: Crb (cerebellum), Ch. plx (choroid plexus), Ctx (cerebral cortex), hyp. th. (hypothalamus), Mb (midbrain), Thal. (thalamus) and wh. mat. (white matter). **(B-C)** Venn diagrams and bar graphs depicting common transcripts and related GO enrichment profiles respectively, in 4 ST annotations with abundant aSyn pathology (pons, midbrain, white matter and choroid plexus, see explanation under Results). **(D)** Representative spatial maps depicting the relative expression of select HALLMARK gene sets (glycolysis, oxidative phosphorylation, fatty acid metabolism, ribosome and mRNA translation machinery, inflammation and circadian rhythm) in cohorts of M83^+/+^ mice.

In contrast, the LS ST profiles in the disease affected regions (pons, midbrain and white matter) reflected significant enrichment of the transcripts belonging to pro-inflammatory, cell proliferative and tissue remodelling processes (inflammatory response, interferon alpha and gamma, complement, TNF signaling via NFkB, G2M checkpoint, mitotic spindle, KRAS signaling)-(Fig. 3A). Of note, annotations encompassing regions with minimal aSyn pathology in the model (cortex, thalamus and hypothalamus) exhibited comparatively unaltered pattern of expression in these pathways (Fig. 3A). Lastly, we found a few unique expression patterns in a subset of pathways, which were predominantly restricted to distinct ST annotations, for instance: E2F targets (cerebellum and midbrain), PI3K-AKT-mTOR signaling (cerebellum and cortex), protein secretion (hypothalamus and midbrain), estrogen response (pons and choroid plexus) and hedgehog signaling (thalamus and cortex)-Fig. 3A.

Then, we selected ST annotation profiles with abundant aSyn aggregation at ES (pons, midbrain, white matter) in the model ^17,18,21^, for further interrogating the common pathways affected during the progression of aSyn pathology. Moreover, the ST data also indicated considerable overlap between the expression patterns in the majority of HALLMARK gene sets in these annotations (Fig. 3A). To these analyses, choroid plexus was also added due to a similar pattern of HALLMARK expression, as well as for the fact that it is a major non-grey matter anatomical element in the ventricular system spanning brainstem ^28^. Despite the relatively high abundance of unique markers (Fig. 2E; *Ttr, Prlr, Ezr, Igfbp2*), it is plausible that the lack of microscopic resolution (inherent to the technology) may underlie the observed overlap in ST expression profiles (Fig. 3A). In these analyses, we found 34 common transcripts with significantly altered expression (log2 FC ≥0.25; adjusted p-value, ≤0.05) across these 4 annotations in association with the ES (Fig. 3B). Gene ontology (GO) enrichment revealed that the common transcripts predominantly belong to the ATP generating metabolic processes (GO:0046034, GO:0009205, GO:0006735, GO:0061718, GO:0006090, GO:0015986, GO:0006119, GO:0072330: *Gapdh, Aldoa, Hspa8, Atp5b, Atp5g1, Spp1, Ndufv1, Pfkm*), axonogenesis (GO:0007409: *Dpysl2, Apoe, Actb, Aplp1, Mbp, Atp5g1*) and synapse organization (GO:0050808: *Apoe, Actb, Hspa8, Sparcl1, Chchd10*).

In contrast, the LS ST profiles were characterized by a larger number (229) of significantly altered across these 4 annotations (Fig. 3C). GO enrichment revealed that the differentially expressed transcripts participate in the regulation of synaptic transmission (GO:0042391, GO:0001505, GO:0007611: *Kif5b, Ppp3ca, Cacnb4, Kcna1, Ank3, Akap9, Rgs7bp, Nrxn1, Scn1a, Pclo, Slc4a4, Gnaq, Rims2, Cacng2, Slc8a1, Gabrb1, Gsk3b, Ckap5, Scn2a, Ank2, Kcnma1, Celf4, Ppp1r9a, Grin2b, Cacnb4, Gpm6b, Slc1a2, Stxbp5l, Nrxn3, Prepl, Braf, Nf1, Dnm1l, Unc13c, Napb, Gad2, Nrxn1, Tmod2, Slc24a2, Map1a, Ube3a, Sgk1, Kmt2a, Cpeb3, Elavl4*), cellular protein sorting and microtubule transport (GO:1903829, GO:0032886, GO:0072698, GO:0010970: *Kif5b, Pcm1, Ccdc88a, Cacnb4, Ank3, Nrxn1, Tpr, Rock2, Myo5a, Bicd1, Sptbn1, Cep290, Map1a, Apc, Tmem30a, Cacng2, Crebrf, Gsk3b, Tcaf1, Dnm1l, Npm1, Map2, Akap9, Macf1, Map1b, Rock1, Taok1, Sgk1, Ckap5, Clip1, Hook3, Golgb1, Cep350, Pura, Dst, Kif1b*) and RNA splicing and metabolism (GO:0008380; GO:1903311: *Malat1, Rock2, Rock1, Tut4, Rbfox1, Rbm25, Mbnl2, Sltm, Srpk2, Cpeb3, Mbnl1, Elavl4, Ncl, Npm1, Celf4, Zbtb7a, Luc7l2, Zfp638, Scaf11, Prpf40a, Tcerg1*).

These and other HALLMARK gene sets can also be visualized as global ST maps using the online interface (URL: https://dreamapp.biomed.au.dk/PD_spatial_mouse_DB/; also see User Guide in Supplementary Information). To illustrate, the ST maps reflect the biphasic pattern of expression (initial upregulation, followed by downregulation) of the pathways controlling glycolysis, oxidative phosphorylation, fatty acid metabolism and mRNA translation machinery (Ribosome)-Fig. 3D; also see Fig. 4-5. Conversely, changes in the expression of transcripts belonging to pathways regulating inflammatory response and circadian rhythm define the LS pathology (Fig. 3D; also see heatmaps in Fig. 5D-E). Apart from the DGE profiles, a number of individual transcripts exhibited a global upward trend, ie., upregulated in ES and further increase in LS (*mt-Nd4l, mt-Atp8, mt-Nd5, mt-Co1, mt-Nd2, mt-Nd3, Spp1, Apod, Foxp2, Slc17a6, Sparcl1, Car2, Gabra1, Rbm3*). Conversely, we also noticed a few transcripts which exhibited a continuous downward trend in the global expression (*Col1a1, Scg2, Lrrc17, Basp1, Bc1, Tatdn1, Tsc22d3, Nr2f2, Nap1l5, Cbx6, Xist, Impact, Hspa5, Gabra2, Syt1, Syt4, Hsp90b1, Ncam1, Gap43, Rps2, Ndn, Rpl9, Cartpt*).

**Figure 4.**
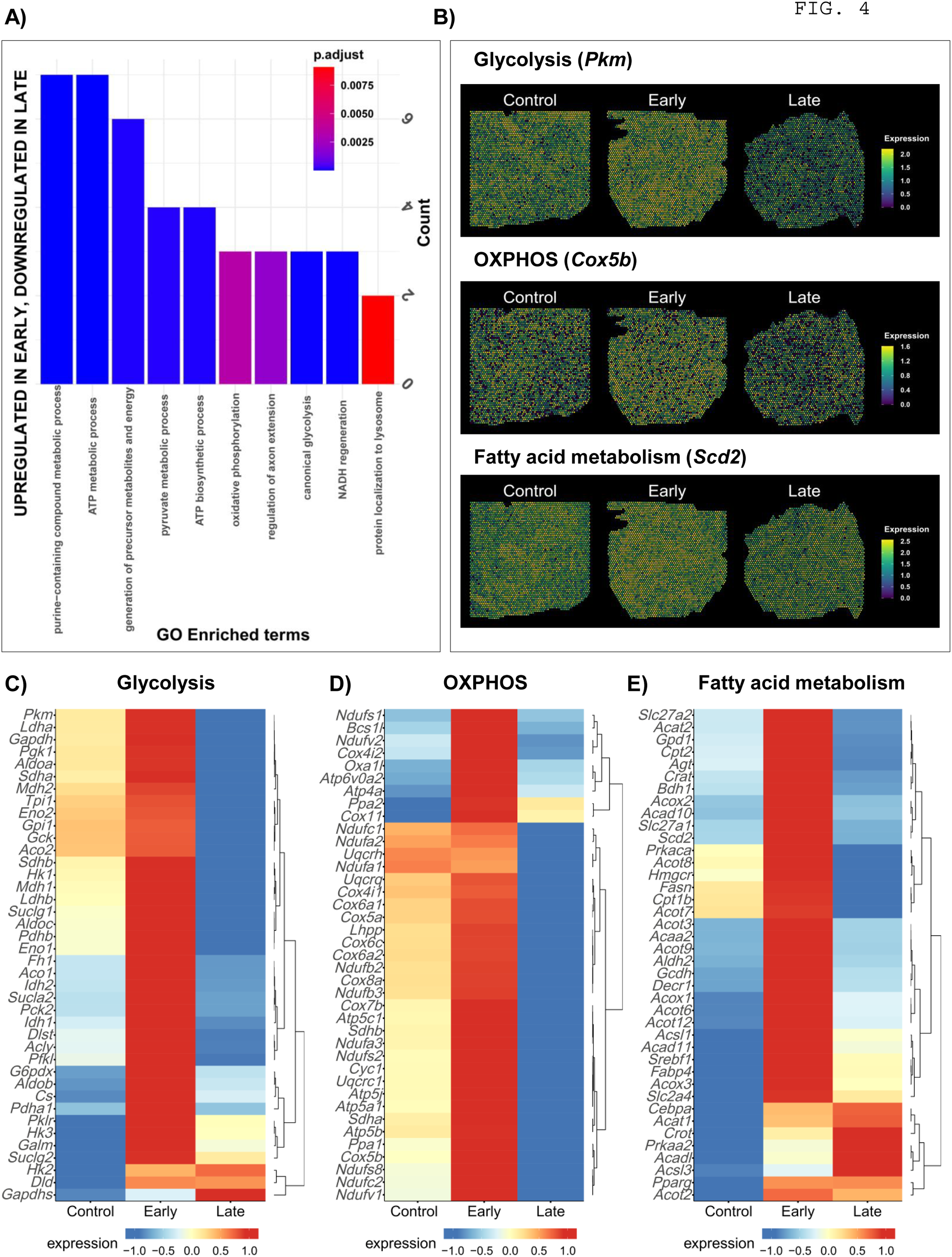
Overview of the differentially expressed gene module/group 3 showing metabolic pathway enrichment in the early stage cohort of M83^+/+^ mice. **(A)** Bar graph depicting GO enrichment profiles across ST annotations in the cohorts of M83^+/+^ mice. **(B)** Representative spatial maps depicting the relative abundance of select transcriptomic markers in metabolic pathways controlling glycolysis (*Pkm*, pyruvate kinase), oxidative phosphorylation (*Cox5b*, subunit 5B of cytochrome c oxidase) and fatty acid metabolism (*Scd2*, stearoyl-CoA desaturase) in sagittal brain sections from cohorts of M83*^+/+^* mice. **(C-E)** Heatmaps depicting the relative expression of select 40 transcripts in metabolic pathways controlling glycolysis/gluconeogenesis, (in C), oxidative phosphorylation (in D) and fatty acid metabolism (in E).- Also see Fig. S1A-C, depicting the complete KEGG pathways shown in Fig. 4C-E.

**Figure 5.**
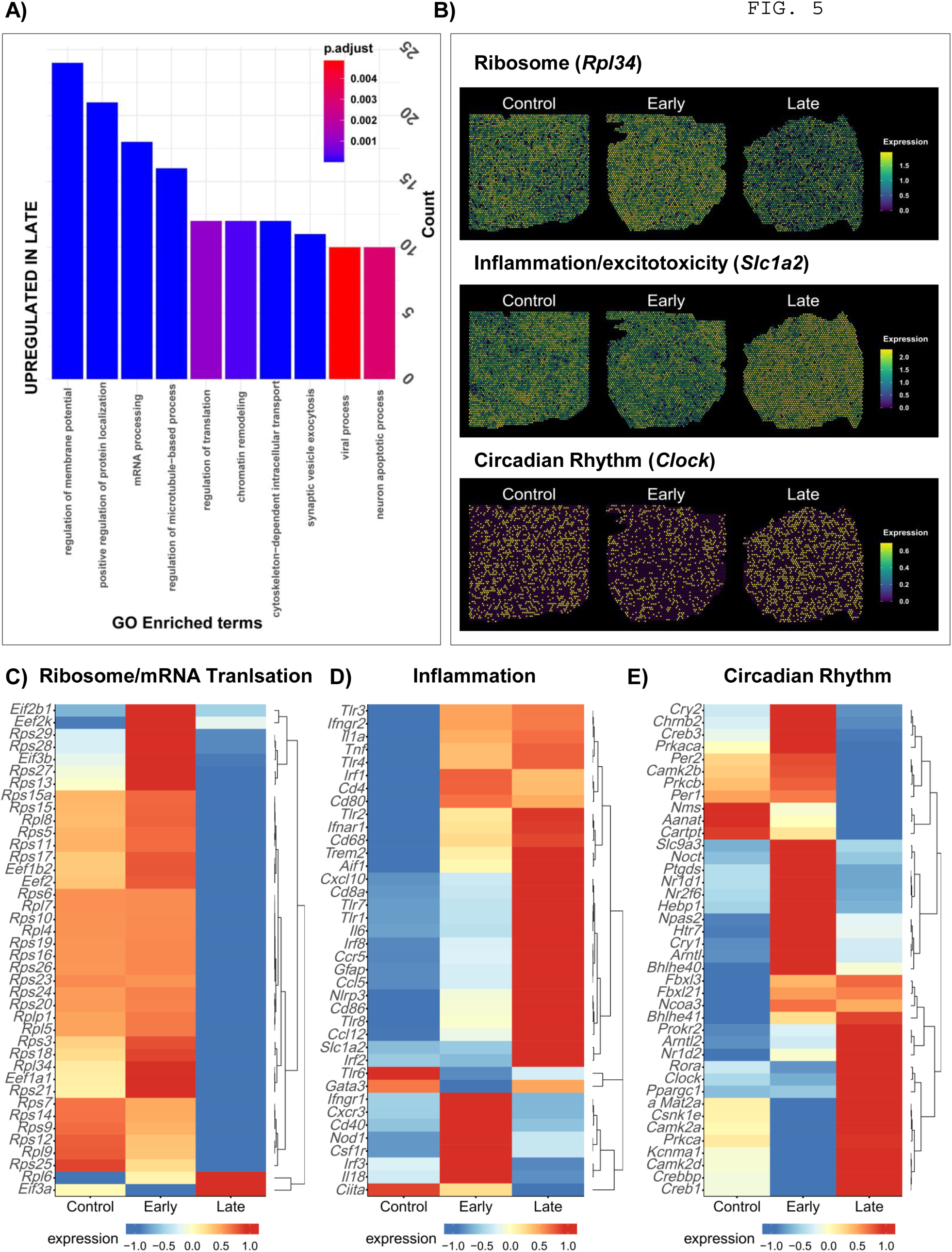
Overview of the differentially expressed gene modules/groups 7 + 8 showing enrichment of pathways controlling mRNA translation, inflammation and circadian rhythm in cohorts of M83^+/+^ mice. **(A)** Bar graph depicting GO enrichment profiles across ST annotations in the cohorts of M83^+/+^ mice. **(B)** Representative spatial maps depicting the relative abundance of select transcriptomic markers in molecular pathways controlling ribosomal assembly (*Rpl34*, a component of large ribosomal subunit), inflammation (*Slc1a2*, encoding the glutamate transporter EEAT2) and circadian rhythm (*Clock*, a master regulator of the pathway) in sagittal brain sections from cohorts of M83*^+/+^* mice. **(C-E)** Heatmaps depicting the relative expression of select 40 transcripts in molecular pathways controlling ribosomal assembly and mRNA translation (in C), immune response and inflammation (in D) and circadian rhythm regulation (in E).- Also see Fig. S2A-C, depicting the complete KEGG pathways shown in Fig. 5C-D.

### The early and late stages of aSyn pathology in the brains of M83^+/+^ mice were defined by differential gene expression modules (DGEMs)

With this global overview of the ST data (Fig. 3), we hypothesized that the DGE profiles associated with ES (prodromal aSyn pathology) potentially reflect a compensatory (tissue homeostatic) phase, while the ST expression profiles seen with LS (advanced aSyn pathology) could underlie the molecular basis of tissue damage in the brain of these rodents. To explore these mechanistic aspects, we performed co-expression analyses to discern DGEMs in the disease affected regions in relation to the stage of aSyn pathology. In these analyses, we discovered 8-9 DGEMs groups/modules based on GO enrichment and the direction of trend in the expression patterns, in comparisons involving ES vs. Control and LS vs. ES. GO enrichment revealed that each of the DGEMs reflects changes in distinct cellular pathways (eg., metabolic processes, neurotransmission, microtubule transport, RNA and DNA metabolism, glial cell differentiation, immune system)-Table S2. To gain further insights into the pathological relevance of these individual DGEMs, we focused on module 3 (upregulated in ES) and modules 7 + 8 (upregulated in LS). First, these DGEMs (modules 3, 7 and 8) recapitulate the global trends in gene expression (Fig. 3A), thus further exploration of these modules was a promising direction for revealing molecular underpinnings of aSyn pathology progression. Second, this focused approach is also commonly used in the precedent ST studies, such that cross-validation of candidate markers in tissue specimen is subsequently made feasible. For instance, Maniatis et al., described 31 ST brain expression modules in a mouse model of motor neuron disease, and then performed validation of a microglial expression program in tissue specimen ^34^.

Hence, we applied an analytical approach incorporating: i) GO enrichment, ii) visualization of ST maps in the mouse brain and iii) heatmaps encompassing curated gene expression profiling using relevant gene sets in the MSigDB (https://bioinf.wehi.edu.au/software/MSigDB/). These analyses are presented in Fig. 4 (for module 3, upregulated in ES) and Fig. 5 (module 7+8, upregulated in LS). As predicted, module 3 reflected the global changes in DGE profiles, dominated by the metabolic pathways, in association with ES (Fig. 4A). Notable examples include: GO processes (GO:0046034, GO:0072521, GO:0006735, GO:0061621, GO:0006091, GO:0006754, GO:0006090, GO:0030516, GO:0006119; *Gapdh, Aldoa, Ndufv1, Atp5b, Pfkm, Atp5g1, Ldhb, Apoe, Dpysl2, Chchd10*; also GO:0061462-protein localization to lysosome: *Cd81, Hspa8*). This is further illustrated in the ST maps for select transcripts in the affected metabolic pathways (Fig. 4B): glycolysis (*Pkm*, encoding pyruvate kinase isoform, also known as the cytosolic thyroid hormone-binding protein), oxidative phosphorylation (*Cox5b*, encoding the subunit 5B of cytochrome c oxidase in the mitochondrial respiratory complex IV) and fatty acid metabolism (*Scd2*, encoding stearoyl-CoA desaturase, involved in the biosynthesis of fatty acids and mitochondrial beta-oxidation). Curated gene expression analyses revealed perturbed expression of several metabolic transcripts in association with ES, as shown for glycolysis and gluconeogenesis (Fig. 4C), oxidative phosphorylation (Fig. 4D) and fatty acid metabolism (Fig. 4E). Moreover, a few transcripts in these pathways exhibited a persistent upregulation in both the ES and LS, for example *Hk2* (encoding a hexokinase isoform, mediating phosphorylation of glucose for glycolysis), *Dld* (encoding dihydrolipoamide dehydrogenase, an oxidoreductase in the gluconeogenesis pathway), *Acat1* (encoding acetyl-CoA acetyltransferase 1, a crucial mediator of the mitochondrial beta-oxidation), and *Pparg* (encoding the peroxisome proliferator-activated receptor gamma-PPAR-γ, an important factor controlling lipid metabolism with a role in inflammatory response). A pictorial overview of the expression pattern in these curated pathways is presented in Fig. S1A-C.

As mentioned above, the ST modules 7 + 8 were uniquely upregulated in the LS and possibly indicate molecular and cellular processes associated with tissue damage consequent to chronic aSyn aggregate pathology. This was corroborated by GO enrichment (in Fig. 5A) revealing perturbed regulation of membrane potential, microtubule transport and synaptic vesicle exocytosis (GO:1903829, GO:0032886, GO:0030705, GO:0042391, GO:0016079: *Map1b, Ank3, Rock2, Nrxn1, Map1a, Tcaf1, Kif5b, Cacng2, Cacnb4, Sptbn1, Ccdc88a, Npm1, Cep290, Bicd1, Tmem30a, Myo5a, Apc, Crebrf, Pcm1, Dnm1l, Clip1, Sgk1, Map2, Rock1, Taok1, Akap9, Ckap5, Macf1, Tpr, Pura, Kif1b, Dst, Hook3, Rgs7bp, Celf4, Ank2, Kcna1, Pclo, Scn2a, Ppp3ca, Rims2, Kcnma1, Gnaq, Scn1a, Ppp1r9a, Slc8a1, Grin2b, Slc4a4, Gabrb1, Napb, Stxbp5l, Prepl, Braf, Unc13c, Nrxn3*), mRNA processing and ribosomal translation (GO:0006397, GO:0006417, GO:0016032: *Malat1, Celf4, Rbm25, Elavl4, Zbtb7a, Srpk2, Cpeb3, Luc7l2, Npm1, Ncl, Mbnl1, Rbfox1, Prpf40a, Scaf11, Mbnl2, Rbm26, Sltm, Tcerg1, Eif5, Eif5b, Secisbp2l, Tut4, Tpr, Rsf1, Eif3a, Bicd1, Mphosph8, Eea1, Ssb, Tasor;* also see ST map in Fig. 5B-top panel, showing the downregulation of *Rpl34* which encodes a component of large ribosomal subunit), neuronal apoptosis (GO:0051402: *Trim2, Scn2a, Gsk3b, Srpk2, Rock1, Npm1, Kcnma1, Nf1, Ralbp1, Braf*) and chromatin remodelling (GO:0006338, GO:0051402: *Meg3, Kcnq1ot1, Atrx, Zbtb7a, Rsf1, Npm1, Pbrm1, Rif1, Mphosph8, Smarca5, Tpr, Tasor, Trim2, Scn2a, Gsk3b, Srpk2, Rock1, Kcnma1, Nf1, Ralbp1, Braf*). Curated gene expression analyses using MSigDB gene sets (in Fig. 5C-E) indicated a profound decline in the transcripts involved in the ribosomal assembly and mRNA translation, including several initiation and elongation factors (Fig. 5C). In line with prior studies, which implicate widespread neuroinflammation at LS in this model ^17–20^, several transcripts belonging to pathways mediating immune regulation, excitotoxicity and inflammatory gliosis were also upregulated (Fig. 5D; exemplified by *Aif1*, encoding allograft inflammatory factor 1/Iba1-a marker of microglia and *Gfap*, encoding glial fibrillary acidic protein-a marker of reactive astrogliosis; also see ST map in Fig. 5B-middle panel, showing the upregulation of *Slc1a2*, which encodes the glutamate transporter EEAT2 involved in sodium dependent clearance of glutamate from synapses). A pictorial overview of the expression pattern in these curated pathways is presented in Fig. S2A-C.

Next, we wanted to investigate if the DEGs in the brains of M83^+/+^ mice are recapitulated in gene expression studies in PD. For this purpose, we performed curated gene expression analyses of patient-derived microarray datasets using the Gene Expression Omnibus (GEO) repository, as described previously ^35^. The datasets and brain regions examined included: 1) GSE7621 (substantia nigra-SN) ^36^, 2) GSE43490 (SN, dorsal motor nucleus of vagus-dmX and locus coeruleus-LC) ^37^, 3) GSE20146 (globus pallidus, interna-GPi;) ^38^ and 4) GSE26927 (SN) ^39^. For these analyses, we selected transcripts in the ST data which were common to the disease affected annotations (pons, midbrain, white matter, at both ES and LS) across all DGEMs, encompassing both the ES (395 transcripts) and LS (831 transcripts). Using cut-off criteria (a log2 FC threshold of ≥±0.25 and an adjusted p-value ≤0.05, in at least 2 PD microarray datasets), the results of these analyses are presented in Fig. S3A-B.

In brief, we discovered several transcripts, which were upregulated both in the ST dataset (M83^+/+^ mice, Fig. S3A) an in PD microarray studies (Fig. S3B; GO:1901655, GO:0033574, GO:0071384: *ANK1, APLP1, BBX, CA2, CREBBP, DCN, EDIL3, GPR37, HIPK2, IGF1R, KCNJ10, MBNL1, MGA, NFIA, NKTR, NR3C1, PLEKHB1, PNISR, ROCK2, RPS6KB1, SGK1, SIRPA, SON, SPOP, SPP1, SRRM2, YBX1, YLPM1, ZDHHC20*; but note *CCK* and *SNCA*). Comparatively, only a few downregulated ST transcripts (Fig. S3A) exhibited similar expression pattern in the PD microarrays examined (Fig. S3B; GO:0043532: *ASIC2, ATP5B, ATP6V1C1, ATPIF1, CADM3, CBLN1, GABARAPL1, GRIN, PINK1, PTPRN, RELL2, TH*). Intriguingly, many more of the ST transcripts, which were downregulated in rodent brain, were found to be upregulated in the PD datasets (GO:0043484, GO:0080090, GO:0019222: *CUTA, DHCR24, EIF4H, ELOVL5, EWSR1, H2AFZ, H3F3A, HNRNPA2B1, HNRNPH1, MID1IP1, NR2F2, PACS2, PTGES3, RNF220, RPL21, RSRP1, SPOCK1, SQSTM1, TMEM50B, TUBB4A, VAMP2, ZMIZ2*). Taken together, these data (Fig. S3A-B) point to distinct molecular signatures associated with aSyn-induced proteopathic stress in the nervous system.

### aSyn pathology in the brains of M83^+/+^ activated distinct transcriptional programmes

Next, we resorted to an upstream analyses approach for revealing potential master regulators, which orchestrate the activity of a set of limited downstream effectors observed in omics datasets. One such approach encompasses the determination of transcription factors (TFs), by analyzing the promoters and enhancers of the DEGs, and constructs TF regulons ^40^. Using a similar strategy, we wanted to determine if the ST profiles (Fig. 3-5) can be explained by variations in specific transcriptional programmes. These analyses revealed that several TF regulons were differentially affected in association with ES or LS aSyn pathology (Fig. S4A: TF regulons UP in ES and DOWN in LS, and Fig. S4B: TF regulons UP in LS). Based on the canonical role of the TFs identified, the transcriptional programmes correspond to metabolic regulation (*Eno1b, Srebf2, Klf7, Neurod1Tcf7l2, Cebpa, Esrra, hnf1b, Nrf1, Rara, Rxra, Rxrb, Rxrg, Srebf1, Srebf2, Thra, Xbp1, Zfp692*), immune response (*Atf7, Cebpg, Ebf1, Elf1, Elf3, Foxp1, Hivep2, Ikzf2, Irf2, Irf8, Nr3c1, Rfx7, Spi1, Stat1*; *Gmeb2, Irf3, Nfatc4, Nr1h3, Rfx1, Rfxank, Sox12, Zfp580*), and circadian rhythm (*Clock, Creb1, Ep300, Klf9, Nr1d2, Prox1, Sp1*; *Crem, Dbp, Mettl3, Nkx2.1, Nr1d1*). In addition, the ST data also indicated modifications in the expression of some TF regulons implicated in the survival of dopaminergic neurons and/or dopamine synthesis (*Phox2a, Pitx3, Zic2, Xbp1)* ^41^. Nevertheless, a substantial number of TF regulons in the dataset correspond to general transcriptional regulation through RNA polymerase II (52 TFs), and cell cycle and differentiation (43 TFs). It is noteworthy that the majority of transcriptional programmes were found to be altered across all ST annotations regardless of the disease burden, with few exceptions (for instance, *Rxra* regulon in pons and *Rara* regulon in cerebellum-Fig. S4A).

Among the several DGE ST profiles in relation to the stage of aSyn pathology (Fig. 4-5), the findings related to the TF regulons controlling circadian rhythm (GO:0042753) were particularly interesting (also see Fig. 3D-HALLMARK ST panel, circadian rhythm). In addition to regulating the hormonal regulation of energy metabolism and sleep-wake cycles in mammals (eg., *Clock*), several of the core circadian TFs are crucial mediators of synaptic plasticity (eg., *Creb*), immune response (eg. *Sp1*) and regulation of mRNA translation (eg., *Mettl3*). Hence, we performed curated gene expression analyses for probing the expression of key transcripts involved in the regulation of circadian rhythm. Not surprisingly, we found distinct transcriptomics signatures of perturbations in these pathways, as illustrated by the upregulation of the master circadian regulator *Clock* and downregulation of the circadian repressors (*Per1, Cry2, Nr1d1*) in association with LS aSyn pathology-Fig. 5E (also see, Fig. 5B-bottom panel, ST map for *Clock*). In addition, the ST data (Fig. 5E) also indicated upregulated expression of *Creb* and its enhancer *Crebbp* (encoding the histone acetyltransferase, CREB-binding protein CREBBP/CBP/KAT3A). Taken together, the data (Fig. 3-5) suggest that the progression of aSyn pathology in the nervous system of this rodent model is a potent trigger for metabolic perturbations and promoting a pro-inflammatory milieu (Fig. 4-5). In turn, these processes lead to aberrant regulation of circadian homeostasis and may underlie clinically relevant phenotypes in models of synucleinopathies (elaborated under Discussion).

### Expression of CREBBP protein was not correlated to the extent of aSyn pathology in the brains of M83^+/+^ mice and in post-mortem PD brains

Then, we wanted to investigate whether the transcriptional response in the ST data suggesting enhanced CREB mediated signaling (Fig. 5E) is also reflected in the expression of proteins regulating this pathway. For this purpose, we performed IF analyses for the detection of CREBBP/CBP, which binds phosphorylated CREB in nucleus and enhances its transcriptional activity ^42^. This crucial regulatory mechanism controls the expression of cAMP-responsive genes, which are involved in large number of processes, including dopaminergic signaling ^43^. In addition, this 265 kDa protein is a known transcriptional coactivator of the circadian clock complex ^44^. CREBBP is ubiquitously expressed in neuronal and glial cells, and its expression has been suggested to decline with ageing and in models of neurodegenerative diseases ^45,46^. Importantly, the choice of pursuing this factor in subsequent analyses was also dictated by the fact that *CREBBP* was consistently upregulated across the PD microarray datasets examined in this study (Fig. S3B).

First, we visualized *Crebbp* ST maps and found a slight, but significant increase in the LS cohort vs. ES (eg., Pons: Log2 FC= 0.19, adjusted p-value= 3.48E-10; Midbrain: Log2 FC= 0.19, adjusted p-value= 9.96E-05; Cerebellum: Log2 FC= 0.25, adjusted p-value= 8.09E-13 and Thalamus: Log2 FC= 0.21, adjusted p-value= 2.92E-05)-Fig. 6A. Not surprisingly, this expression pattern in the regions is reflective of the relative abundance of the *Crebbp* transcript in the mouse brain (Allen Mouse Brain Atlas, in-situ hybridization). Next, we performed IF detection of CREBBP in the sagittal brain sections from M83^+/+^ cohorts, with focus on brain regions with abundant aSyn pathology (GRN in pons, PAG in midbrain) and brain regions affected to a mild degree in this model (DCN in cerebellum, MD thalamus) ^18,20^-also see Fig. 1C. Our analyses indicated an overall trend of progressive increase in CREBBP IF positive cells in all the regions examined, regardless of the extent of aSyn pathology (Fig. 6B-C). Albeit, a clear difference could be discerned in the LS brains compared to the controls or ES samples (Fig. 6B-C). Due to the small samples size, the data did not reach statistical significance. Intriguingly, the largest effect was seen in the mediodorsal (MD) thalamic nuclei (Fig. 6C; compare DM to GRN, PAG and DCN), an area with sparse aSyn pathology in the model (Fig. 1C). Although, the IF results (Fig. 6B-C) corroborate the findings in ST data (ie., LS aSyn pathology is associated with elevated CREBBP expression-Fig. 5E), the lack of correlation to the relative abundance of aSyn pathology in different regions (Fig. 1B-C) is perplexing.

**Figure 6.**
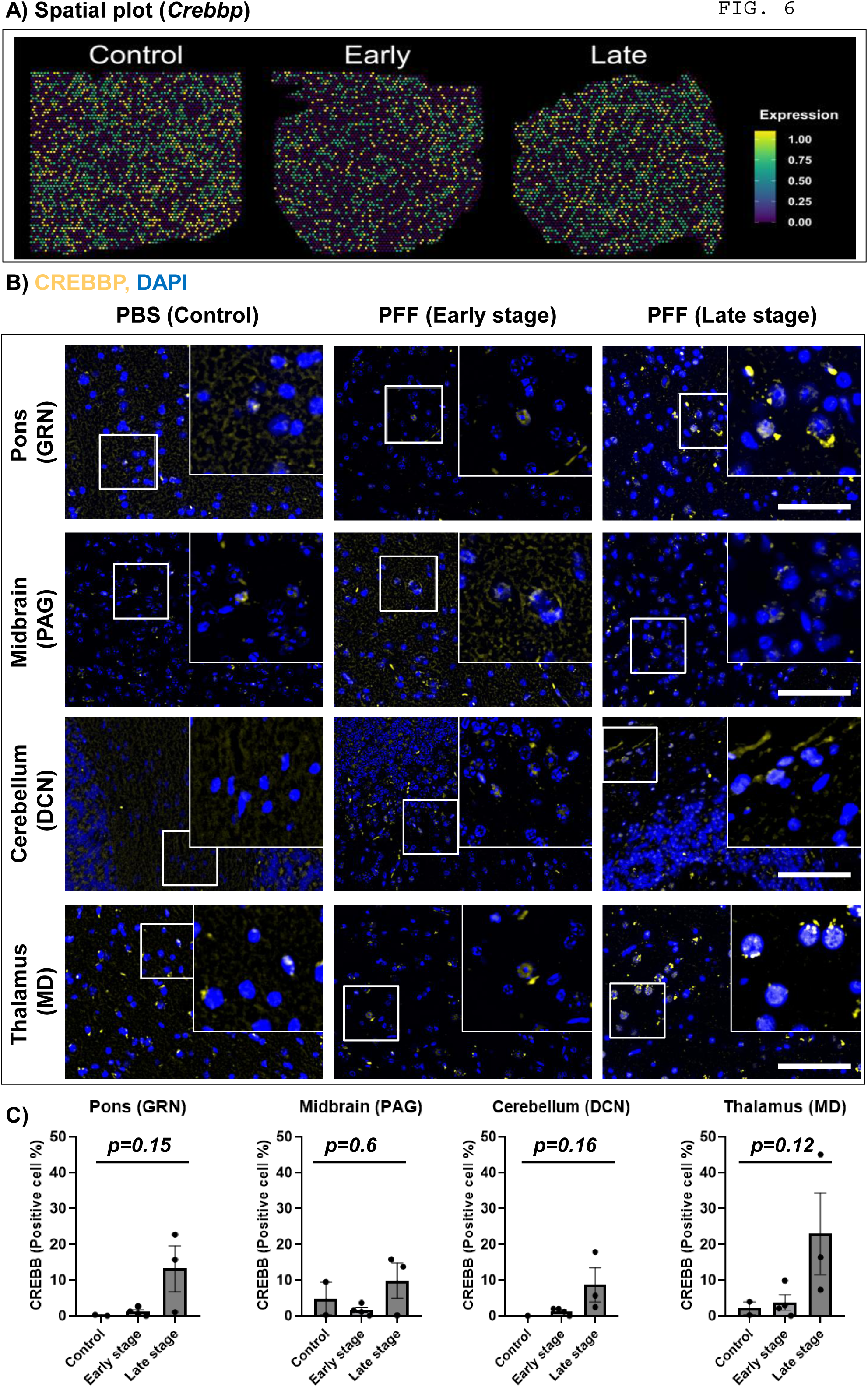
ST maps and IF detection of CREBBP/CBP in brains of M83^+\+^ mice. **(A)** Representative spatial maps depicting the relative abundance of *Crebbp* in sagittal brain sections from cohorts of M83*^+/+^* mice. **(B)** Representative (20X) images showing CREBBP/CBP IF in the gigantocellular nuclei (GRN, in pons), periaqueductal grey (PAG, in midbrain), deep cerebellar nuclei (DCN, in cerebellum) and mediodorsal nuclei (MD, in thalamus). The inset show 63X magnified views from the regions. Scale bar= 50 µm. **(C)** Bar graphs depicting quantification of CREBBP/CBP IF in GRN, PAG, DCN and DM of the experimental cohorts, as indicated. Error bars depict Mean IF intensity ± SD as % of cells in total area (PBS, n=2; DPI-45, n=3 and DPI-75, n=3; See Methods). Statistics in Fig. 5C: Kruskal-Wallis ANOVA (p-values on graphs), Dunn’s multiple comparisons (non-significant).

For comparison, we performed IF detection of ROCK2 (Rho Associated Coiled-Coil Containing Protein Kinase 2) in the brains of M83^+\+^ mice, another disease-relevant pathway identified in the ST data (Fig. S3A, *Rock2*) and in PD microarrays (Fig. S3B, *ROCK2*). This pleiotropic serine/threonine kinase is expressed throughout the neuraxis, and is the predominant isoform (compared to ROCK1) in the nervous tissue ^47^. In response to activation by Rho GTPases, ROCK2 phosphorylates several cellular targets involved in actin-mediated cytoskeletal organization, cytokinesis, and neurite stability ^47,48^. Furthermore, increased expression (neuronal and glial) of ROCK isoforms has been reported within the disease affected regions in AD, PD and atypical parkinsonism ^48–50^. Our IF analyses indicated substantially enhanced expression and/or localization of ROCK2 in brain regions with progressive aSyn pathology in the rodent model (Fig. S5A-C; compare GRN, PAG and DCN to DM; also see Fig. 1C).

Lastly, we resorted to perform the immunodetection of CREBBP in PD post-mortem brain, since we were encouraged by the data indicating enhanced CREBBP expression in association with LS pathology in M83^+/+^ mice (Fig. 6B-C), as well as *CREBBP* upregulation in *CREBBP* across the PD microarray datasets (Fig. S3A, *CREBBP*). Using p-aSyn (S129) IHC in midbrain sections from control and PD cases (Table S1), we evaluated the extent of aSyn aggregate pathology in SN and PAG regions ^5,30–32^. As expected, these analyses revealed significant p-aSyn (S129) immunopositive cells in PD midbrain, within both the SN (controls: 0.36 ± 0.05; PD: 4.02 ± 0.63, p-value= 0.002; n=6/group; cells/mm^2^) and the PAG (controls: 0.29 ± 0.1; PD: 3.86 ± 0.44, p-value= 0.004; n=5-6/group). In serial midbrain sections, we performed IHC analyses for CREBBP in these regions, which suggested enhanced CREBBP expression in PD, but did not reach statistical significance. In summary, we detected a 3-4 fold increase in nuclear CREBBP immunopositve cells, in both the SN (controls: 3.46 ± 1.35; PD: 9.22 ± 4.17, p-value= 0.69; n=6/group) and the PAG (controls: 1.28 ± 0.48; PD: 4.33 ± 2.23, p-value= 0.99; n=5-6/group). However, a closer examination of the IHC data within each control and PD case suggested that aSyn pathology (p-aSyn, S129) and CREBBP immunopositivity were not strongly correlated in the SN (Spearman’s correlation coefficient: controls, 0.13; PD: 0.14, n=6/group). In contrast, a moderate degree of correlation between the two markers could be deduced in the PAG (Spearman’s correlation coefficient: controls, 0.65; PD: 0.50, n=5-6/group). Among the 6 PD cases examined, the highest CREBBP IF was detected in midbrain (SN and PAG) of PD-3 and PD-4 (Fig. S6A-B), with the clinical diagnosis of PD compounded by dementia (Table S1). Although the data suggested enrichment of CREBBP protein with the progression of aSyn pathology in brains of M83^+/+^ mice (Fig. 6B-C), a degree of mismatch in PD midbrain sections (Fig. 7A-B, Fig. S6A-B) precludes major conclusions.

**Figure 7.**
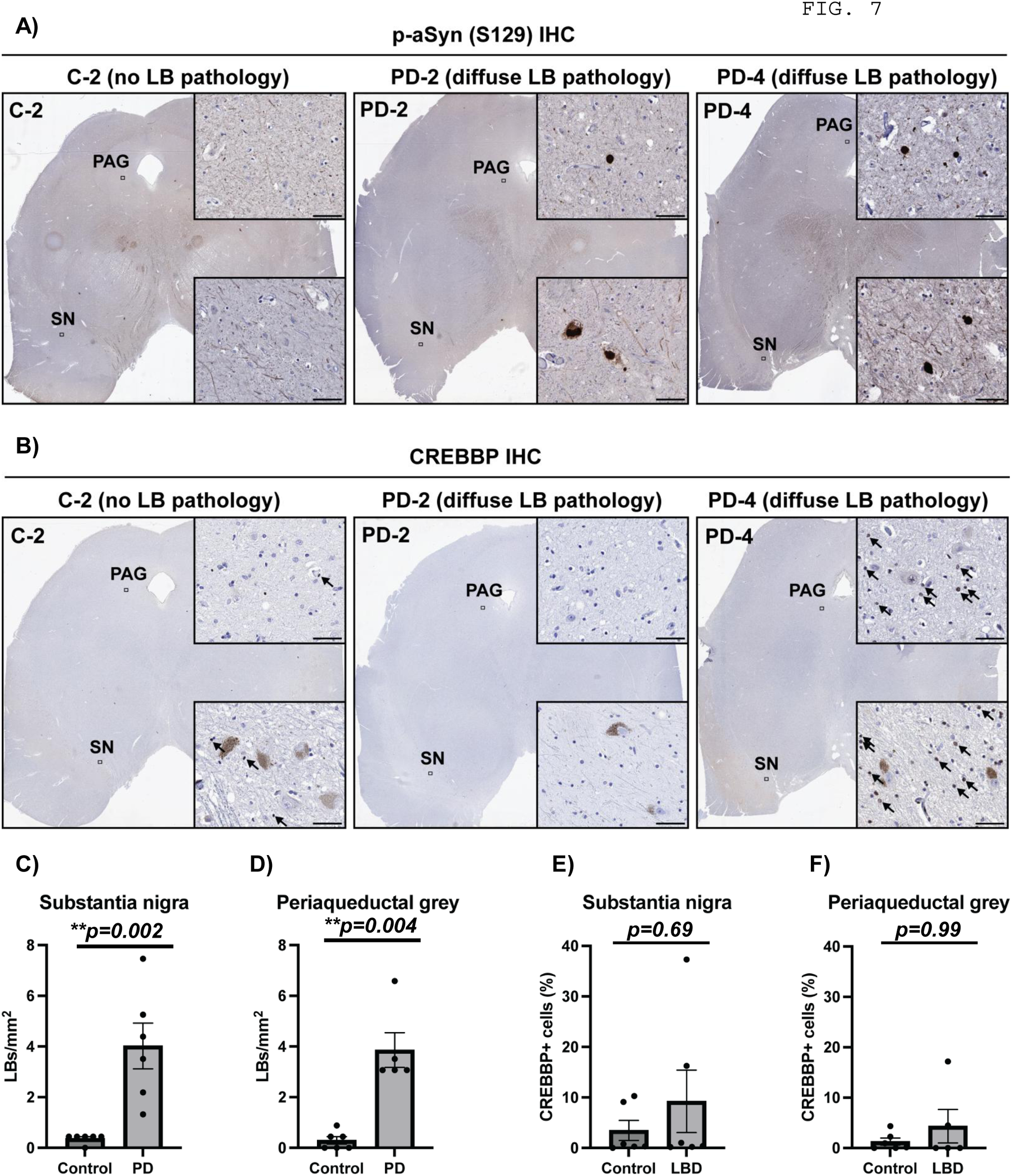
Immunostaining of phospho-alpha synuclein (p-aSyn, S129) and CREBBP/CBP in post-mortem human brain sections. **(A-B)** Representative images showing IHC detection of p-aSyn (S129; in A) and CREBBP/CBP in (B) in post-mortem midbrain sections from controls and PD cases (Table S1). Insets show high magnification images from the substantia nigra (SN) and periaqueductal grey (PAG), reflecting prominent Lewy-related aSyn pathology in PD and visible lack of staining in the controls. Scale bar in insets = 50 µm). **(C-D)** Bar graphs depicting quantification of Lewy body pathology (p-aSyn, S129) in the SN (in C) and PAG (in D). Graphs display Mean ± SEM of Lewy bodies per mm^2^. **(E-F)** Bar graphs depicting quantification of CREBBP/CBP IHC in the SN (in E) and PAG (in F), expressed as Mean % ± SEM of immunopositive cells in the region (see Methods). Statistics in Fig. 7C-F: Pair-wise comparisons using Mann-Whitney test (p-values on graphs).

## DISCUSSION

Circumstantial evidence, gleaned from studies in neuronal cultures and animal models, implicates that the process of aSyn aggregation is a potent stress factor that disrupts synaptic function and impairs neuronal survival in neurodegenerative proteinopathies, including PD ^1,4,7,51^. Therefore, preventing aSyn aggregation and/or therapeutically modulating cellular pathways of aSyn neurotoxicity represent highly desired pursuits in clinical drug development for PD and related diseases ^52^. Moreover, there is growing support for the notion that pathological forms of aSyn likely propagate in a prion-like fashion, such that there is progressive damage to multiple neuronal populations (in addition to the dopaminegric neurons in SN*pc*), which in turn is potentially linked to the heterogeneity in clinical presentation ^5,8,11,53^. In this regard, there are suggestions that pathological aSyn deposition in the nervous system begins several years prior to the onset of motor disability seen in clinical parkinsonism ^1,2,5,11,22^. Accordingly, the molecular determinants of the resilience and/or vulnerability of distinct neuronal populations to incipient aSyn pathology, which in turn dictate the patterns of neuronal dysfunction leading to PD symptoms, are increasingly being explored ^54–57^.

In the present study, we applied ST to brain sections collected from a mouse model of extra-nigral synucleinopathy in brainstem ^15,18,19^, to interrogate: i) the cellular adaptations in the brain during the prodromal phase (ES) of synucleinopathy, which are potentially counteracting the neurotoxicity of aSyn (hence, the relative lack of gross motor phenotypes) ^18^, and ii) perturbations in the cellular milieu indicating progressive cellular aSyn pathology, which herald the onset of symptomatic phase ^15,18,19^. Our ST data support a mechanistic model (Graphical Abstract) which supports the hypothesis that progressive aSyn pathology triggers a compensatory global transcriptional response affecting metabolic pathways (module 3; Fig. 4), which is a harbinger of impending energy crisis and culminates in defective mRNA translation and neurotransmission (module 7 + 8; Fig. 5). This may potentially explain the phenotypic progression from ES to LS in the model, such that the deleterious effects of incipient ES pathology on neuronal function are masked by local metabolic adaptations, which eventually fail with the progression of the disease. Below, we discuss some of the salient takeaways from the present study, first with examples of concrete biological factors, followed by some key relevant examples in the published literature.

The most striking observation from the present study was that the incipient aSyn pathology in brainstem triggers the expression of molecular mediators reflecting enhanced energy flux in mitochondrial metabolic pathways (Fig. 3-5). At first glance, these observations appear counterintuitive to the prevailing notion, which implicates mitochondrial deficiency and impaired mitochondrial respiration as critical drivers of neuronal loss during the pathogenesis of PD ^58,59^. This notion is also supported by the fact that direct application of mitochondrial toxins (e,g. 1-methyl-4-phenyl-1,2,3,6-tetrahydropyridine-MPTP, rotenone, 6-hydroxydopamine) is a highly reproducible method for inducing PD-like dopaminergic cell loss in experimental models ^51,60^. Moreover, several microarray bulk-RNAseq and single-cell RNAseq studies also reinforce the etiological relevance of mitochondrial dysfunction to progressive neuropathology in PD (examples in ^61–63^, also see detailed review of additional studies including limitations ^64^). Furthermore, rare genetic defects in cellular pathways involved in healthy mitochondrial function and integrity cause familial forms of parkinsonism. In this regard, the genetic association of missense variants of *PRKN* (encoding Parkin, an E3-ligase), *PINK1* (encoding the PTEN-induced putative kinase) and *PARK7* (encoding DJ1-, protein deglycase) to autosomal recessive PD is well-established ^1,2,59^. Related to this, our ST data in M83^+/+^ mice suggest perturbed expression of Gpr37, a substrate for Parkin, in relation to aSyn pathology (Fig. S3A, *Gpr37*), and was corroborated in the PD microarray datasets (Fig. S3B, *GPR37*). This transcript encodes an orphan G protein-coupled receptor with putative role in negatively regulating myelination, and is reported to accumulate in misfolded states within the PD brain ^65^. Moreover, the ST data indicate that the progression of aSyn pathology in the brains of M83^+/+^ mice eventually culminates in transcriptional perturbations leading to mitochondrial deficiency (Fig. 3-4, Fig. S1), as has been suggested in the context of neurodegeneration in PD ^58,59^. Interestingly, a recent study in amyloid-precursor protein (APP) knock-in mice has reported similar biphasic transcriptomic alterations in mitochondrial metabolic pathways, in relation to progressive neurodegenerative amyloid pathology *in vivo* ^66^.

The second remarkable finding in the ST data in relation to advanced aSyn pathology hints towards defects in ribosomal mRNA translation machinery (Fig. 4B-C). The pathogenic relevance of this aspect is highlighted by studies showing reduced expression of several initiation and elongation factors in neurodegenerative diseases, including PD (reviewed elsewhere ^67^). Moreover, several reports indicate hyperphosphorylation of the eukaryotic translation initiation factor-2 (eIF2α) ^68^, eukaryotic elongation factor-2 (eEF2) ^25^, AMP-responsive kinase (AMPK) ^69^ and mTOR and its downstream signaling targets in PD and related neurodegenerative conditions ^70^. The expected net effect of these biochemical changes in the homeostatic regulation of mRNA translation machinery is a decline in neuronal protein synthesis, an energy consuming process, and activation of the integrated stress response ^67,70^. In neuronal cells, there is considerable evidence that mRNA translation takes place in dendrites and is critical for synaptic plasticity and reorganization. The delivery of synaptic mRNAs, components of translational machinery and synaptic vesicles away from the neuronal cell body requires an intact microtubule transport system, which is also an energy demanding process ^71^. Thus defects in metabolic fuel utilization, as would occur under mitochondrial deficiency, hamper these homeostatic processes and serve as stress signals. Supporting this notion, the ST data show that the expression of *Spp1* (encoding osteopontin, a stress factor involved in cell-matrix interactions and tissue remodeling) is increased in the brains of M83*^+/+^* mice during the progression of aSyn pathology (Fig. S3A). Moreover, the pathogenic relevance of this pathway is also reinforced by the findings in PD microarray datasets (Fig. S3B, *SPP1*), and studies showing increased osteopontin levels in the cerebrospinal fluid and serum in some cohorts of PD patients ^72^. Related to this, a crucial effector mechanism in tissue damage associated with neurodegenerative aggregate pathology involves the activation of ROCK2 (a serine/threonine kinase). Growth inhibitory signals in neuronal milieu are potent activators of RhoA/ROCK2 pathway, which in turn leads to growth cone collapse and axonal degeneration (for instance, through phosphorylation of LIM domain kinase) ^73^. Increased ROCK2 expression and/or markers of RhoA/ROCK2 activity are linked to several neurodegenerative conditions, including PD and atypical parkinsonism ^49,50^ (also see, Fig. S3B, *ROCK2*). Not surprisingly, our ST data also corroborate the pathological relevance of ROCK2 in brains of M83^+\+^ mice, as demonstrated by increased *Rock2* transcript enrichment (Fig. S3A), and IF detection of the protein in association with LS aSyn pathology (Fig. S5B-C).

While the above findings highlight defective homeostatic regulation of energy balance in response to progressive aSyn pathology, it is difficult to pinpoint their role in the etiology of specific PD symptomatology. It is increasingly being recognized that non-motor symptoms (sleep disturbances, pain, olfaction and autonomic dysfunction) are an important cause of reduced quality of life in PD ^1,2,74^, and could arise due to disturbances in circadian regulation ^75,76^. The molecular and genetic evidence to support this notion in clinical studies and animal models is only recently beginning to emerge (reviewed elsewhere ^76,77^). In this regard, our findings related to perturbed expression of transcripts in the pathway regulating circadian rhythm (Fig. 5E) and CREBBP/CBP protein (Fig. 6B-C) are potentially relevant. Cellular studies show that CREBBP/CBP is direct activator of circadian complex CLOCK/BMAL1 (Basic Helix-Loop-Helix ARNT Like 1) in nucleus, which then binds to enhancer elements in promoter regions of core clock gene Period 1 (*PER1*) ^44^. This mechanism seems to be independent of transcriptional activation of CREB by CREBBP/CBP in the canonical regulation of c-AMP responsive gene expression. Our analyses in post-mortem PD brain sections did not conclusively establish a significant correlation between the extent of aSyn pathology and CREBBP/CBP abundance in SN or PAG (Fig. 7A-B). Nevertheless, the data from 2 cases with clinical presentation of advanced PD and dementia (Table S1, PD-3 and PD-4; Fig. S6A-B) are encouraging and warrant further investigations, since defective CREB signaling is implicated in Alzheimer-type neurodegeneration ^45,46^. It is noteworthy that altered sleep-wake cycles and circadian rhythm have been reported in a limited number of studies in rodent models of synucleinopathy ^78–80^. In this context, we have reported that aSyn aggregate pathology in GRN and PAG of PFF aSyn-injected heterozygous M83+/− mice leads to an early onset phenotype, characterized by progressive decline in nocturnal activity in home cage (albeit, these findings are not yet peer reviewed) ^81^.

Lastly, a considerable limitation of the present study,-apart from the small sample size-, is that the major findings are derived from a transgenic rodent model, with PFF-mediated induction of aSyn pathology in the absence of PD-like neurodegeneration (especially in SN*pc*) ^15,18,19^. This limitation is not unique to this model, as recapitulating neuropathological features relevant to PD in association with PD-related phenotypes (motor and non-motor) remains a challenging task ^7,51^. Nevertheless, precedent literature show that this approach in applying ST for molecular investigations in models of neurodegeneration, complemented with limited validation in precious human post-mortem tissue specimen, is both informative and cost-effective ^34,82–84^. For example, Goralski et al., ^83^, applied ST for studying the molecular signatures of cortical Lewy body (LB) aSyn pathology in human post-mortem brain sections (PD, PD with dementia, and dementia with LB), and in brains of wild type mice following PFF aSyn delivery in the striatum. Their analyses revealed several commonly affected pathways (in human brain sections, and in rodent brain 3 months after PFF aSyn injections), with downregulation of mitochondrial metabolic pathways, synaptic markers and unfolded protein response. In comparison, their data indicated upregulation in pathways controlling DNA/RNA integrity and complement factors ^83^. These findings recapitulate features associated with LS pathology in our M83*^+/+^* ST dataset (Fig. 3, Fig. 5 and Fig. S3A), which are also supported by analyses involving PD microarrays (Fig. S3B). Hence, our findings regarding the initial surge in the expression of molecular indicators of metabolic flux in mitochondria (Fig. 3) are intriguing, and worth investigating in relation to the stages of PD pathology. We anticipate that such analyses will be highly informative for biomarker discovery in PD and related synucleinopathies. Lastly, we consider that our ST data will be a valuable aid in transcriptomics-guided drug discovery and repurposing, as exemplified by translational studies using midbrain neurons derived from human-induced pluripotent stem cells from PD patients ^85^ and rodent models of tauopathy ^84^.

## Conclusion

Within the context of complex etiology of idiopathic PD, pathological aSyn aggregation is a potent exacerbating factor. In the present study, we describe the molecular signatures associated with progression of PD-like aSyn pathology in nervous system. Our findings support a mechanistic model whereby the failure of mitochondrial metabolism, following an initial compensatory phase, heralds the onset of transcriptional reprogramming potentially linked to circadian rhythm disturbances in PD.

## Supporting information

Supplemenatry Information Lin et al.

## Acknowledgements

The authors would like to thank Trine Mikkelsen (JRN lab) for help and assistance with the preparation for mouse brain sections for the St study.

## Author Contributions

LL, AD and AJ designed research; LL, NMJ, SAF, AD, DGM, MR and AJ performed research; LL performed bioinformatics analyses and designed the online visualization platform; NMJ, SAF, OAA and AJ analyzed data; IRM provided human post-mortem tissue and contributed to discussion; PHS, MRR, CBV, PHJ and JRN contributed with infrastructural and personnel support; LL, NMJ and AJ wrote the manuscript. All the authors read and approved the manuscript.

---

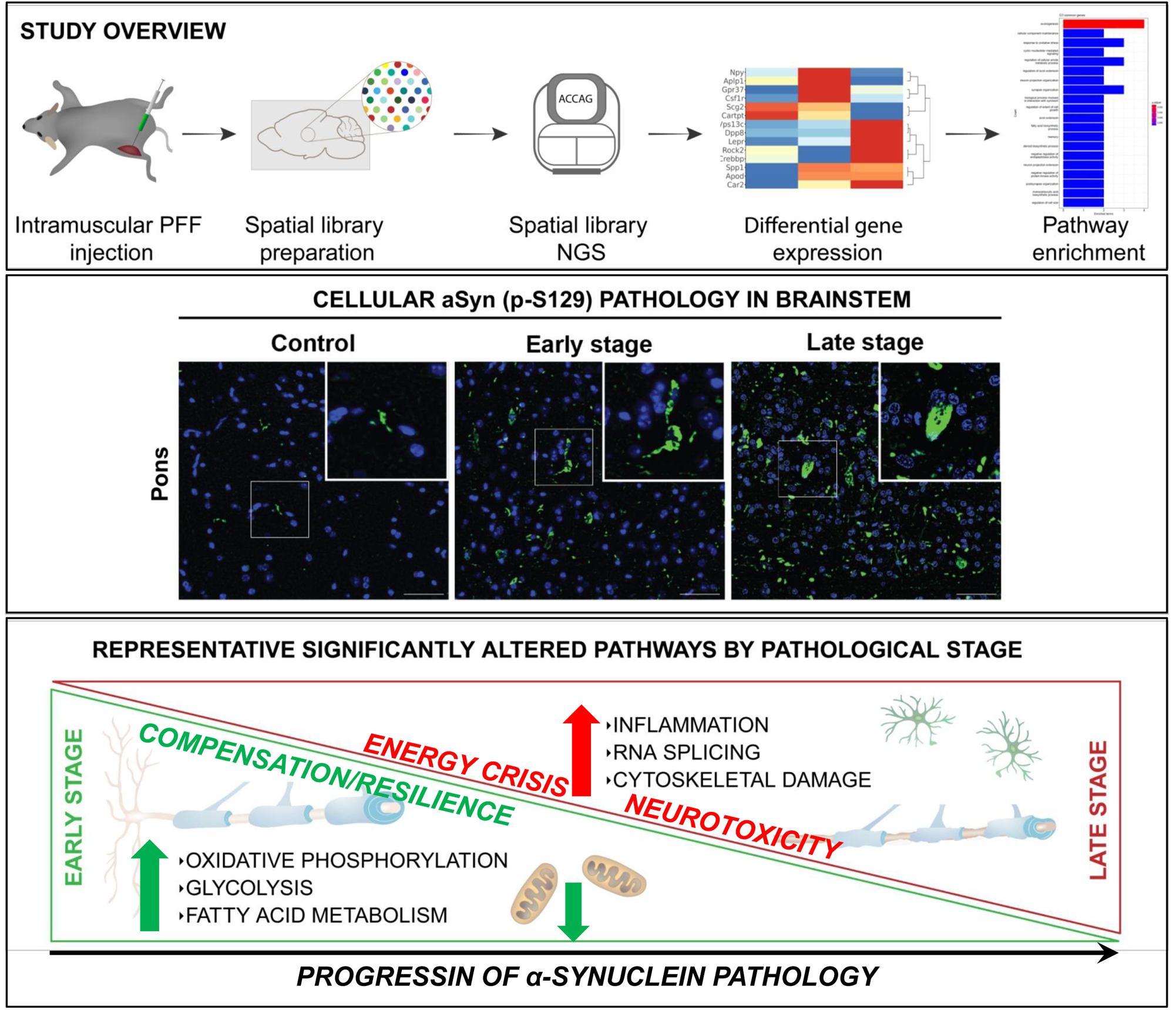

## REFERENCES

1. Kalia, L.V., and Lang, A.E. (2015). Parkinson’s disease. Lancet 386, 896–912. S0140-6736(14)61393-3 [pii] 10.1016/S0140-6736(14)61393-3.

2. Poewe, W., Seppi, K., Tanner, C.M., Halliday, G.M., Brundin, P., Volkmann, J., Schrag, A.E., and Lang, A.E. (2017). Parkinson disease. Nat Rev Dis Primers 3, 17013. nrdp201713 [pii] 10.1038/nrdp.2017.13.

3. Lashuel, H.A., Overk, C.R., Oueslati, A., and Masliah, E. (2013). The many faces of alpha-synuclein: from structure and toxicity to therapeutic target. Nat Rev Neurosci 14, 38–48. 10.1038/nrn3406.

4. Wong, Y.C., and Krainc, D. (2017). alpha-synuclein toxicity in neurodegeneration: mechanism and therapeutic strategies. Nat Med 23, 1–13. 10.1038/nm.4269.

5. McCann, H., Stevens, C.H., Cartwright, H., and Halliday, G.M. (2014). alpha-Synucleinopathy phenotypes. Parkinsonism Relat Disord 20 Suppl 1, S62–67. 10.1016/S1353-8020(13)70017-8.

6. Aniszewska, A., Bergstrom, J., Ingelsson, M., and Ekmark-Lewen, S. (2022). Modeling Parkinson’s disease-related symptoms in alpha-synuclein overexpressing mice. Brain Behav 12, e2628. 10.1002/brb3.2628.

7. Koprich, J.B., Kalia, L.V., and Brotchie, J.M. (2017). Animal models of alpha-synucleinopathy for Parkinson disease drug development. Nat Rev Neurosci 18, 515–529. 10.1038/nrn.2017.75.

8. Borghammer, P. (2021). The alpha-Synuclein Origin and Connectome Model (SOC Model) of Parkinson’s Disease: Explaining Motor Asymmetry, Non-Motor Phenotypes, and Cognitive Decline. J Parkinsons Dis 11, 455–474. 10.3233/JPD-202481.

9. Braak, H., Del Tredici, K., Rub, U., de Vos, R.A., Jansen Steur, E.N., and Braak, E. (2003). Staging of brain pathology related to sporadic Parkinson’s disease. Neurobiol Aging 24, 197–211. S0197458002000659 [pii].

10. Braak, H., Rub, U., Sandmann-Keil, D., Gai, W.P., de Vos, R.A., Jansen Steur, E.N., Arai, K., and Braak, E. (2000). Parkinson’s disease: affection of brain stem nuclei controlling premotor and motor neurons of the somatomotor system. Acta Neuropathol 99, 489–495. 10.1007/s004010051150.

11. Jellinger, K.A. (2019). Is Braak staging valid for all types of Parkinson’s disease? J Neural Transm (Vienna) 126, 423–431. 10.1007/s00702-018-1898-9.

12. Menozzi, E., Macnaughtan, J., and Schapira, A.H.V. (2021). The gut-brain axis and Parkinson disease: clinical and pathogenetic relevance. Ann Med 53, 611–625. 10.1080/07853890.2021.1890330.

13. Van Den Berge, N., Ferreira, N., Mikkelsen, T.W., Alstrup, A.K.O., Tamguney, G., Karlsson, P., Terkelsen, A.J., Nyengaard, J.R., Jensen, P.H., and Borghammer, P. (2021). Ageing promotes pathological alpha-synuclein propagation and autonomic dysfunction in wild-type rats. Brain 144, 1853–1868. 10.1093/brain/awab061.

14. Uemura, N., Yagi, H., Uemura, M.T., Hatanaka, Y., Yamakado, H., and Takahashi, R. (2018). Inoculation of alpha-synuclein preformed fibrils into the mouse gastrointestinal tract induces Lewy body-like aggregates in the brainstem via the vagus nerve. Mol Neurodegener 13, 21. 10.1186/s13024-018-0257-5.

15. Giasson, B.I., Duda, J.E., Quinn, S.M., Zhang, B., Trojanowski, J.Q., and Lee, V.M. (2002). Neuronal alpha-synucleinopathy with severe movement disorder in mice expressing A53T human alpha-synuclein. Neuron 34, 521–533. 10.1016/s0896-6273(02)00682-7.

16. Ayers, J.I., Brooks, M.M., Rutherford, N.J., Howard, J.K., Sorrentino, Z.A., Riffe, C.J., and Giasson, B.I. (2017). Robust Central Nervous System Pathology in Transgenic Mice following Peripheral Injection of alpha-Synuclein Fibrils. J Virol 91. 10.1128/JVI.02095-16.

17. Ferreira, N., Goncalves, N.P., Jan, A., Jensen, N.M., van der Laan, A., Mohseni, S., Vaegter, C.B., and Jensen, P.H. (2021). Trans-synaptic spreading of alpha-synuclein pathology through sensory afferents leads to sensory nerve degeneration and neuropathic pain. Acta Neuropathol Commun 9, 31. 10.1186/s40478-021-01131-8.

18. Ferreira, N., Richner, M., van der Laan, A., Bergholdt Jul Christiansen, I., Vaegter, C.B., Nyengaard, J.R., Halliday, G.M., Weiss, J., Giasson, B.I., Mackenzie, I.R., et al. (2021). Prodromal neuroinvasion of pathological alpha-synuclein in brainstem reticular nuclei and white matter lesions in a model of alpha-synucleinopathy. Brain Commun 3, fcab104. 10.1093/braincomms/fcab104.

19. Sacino, A.N., Brooks, M., Thomas, M.A., McKinney, A.B., Lee, S., Regenhardt, R.W., McGarvey, N.H., Ayers, J.I., Notterpek, L., Borchelt, D.R., et al. (2014). Intramuscular injection of alpha-synuclein induces CNS alpha-synuclein pathology and a rapid-onset motor phenotype in transgenic mice. Proc Natl Acad Sci U S A 111, 10732–10737. 10.1073/pnas.1321785111.

20. Sorrentino, Z.A., Xia, Y., Funk, C., Riffe, C.J., Rutherford, N.J., Ceballos Diaz, C., Sacino, A.N., Price, N.D., Golde, T.E., Giasson, B.I., and Chakrabarty, P. (2018). Motor neuron loss and neuroinflammation in a model of alpha-synuclein-induced neurodegeneration. Neurobiol Dis 120, 98–106. 10.1016/j.nbd.2018.09.005.

21. Ayers, J.I., Riffe, C.J., Sorrentino, Z.A., Diamond, J., Fagerli, E., Brooks, M., Galaleldeen, A., Hart, P.J., and Giasson, B.I. (2018). Localized Induction of Wild-Type and Mutant Alpha-Synuclein Aggregation Reveals Propagation along Neuroanatomical Tracts. J Virol 92. 10.1128/JVI.00586-18.

22. Seidel, K., Mahlke, J., Siswanto, S., Kruger, R., Heinsen, H., Auburger, G., Bouzrou, M., Grinberg, L.T., Wicht, H., Korf, H.W., et al. (2015). The brainstem pathologies of Parkinson’s disease and dementia with Lewy bodies. Brain Pathol 25, 121–135. 10.1111/bpa.12168.

23. Chu, W.T., DeSimone, J.C., Riffe, C.J., Liu, H., Chakrabarty, P., Giasson, B.I., Vedam-Mai, V., and Vaillancourt, D.E. (2020). alpha-Synuclein Induces Progressive Changes in Brain Microstructure and Sensory-Evoked Brain Function That Precedes Locomotor Decline. J Neurosci 40, 6649–6659. 10.1523/JNEUROSCI.0189-20.2020.

24. Delaidelli, A., Richner, M., Jiang, L., van der Laan, A., Bergholdt Jul Christiansen, I., Ferreira, N., Nyengaard, J.R., Vaegter, C.B., Jensen, P.H., Mackenzie, I.R., et al. (2021). Alpha-Synuclein pathology in Parkinson disease activates homeostatic NRF2 anti-oxidant response. Acta Neuropathologica Communications 9, 105. 10.1186/s40478-021-01209-3.

25. Jan, A., Jansonius, B., Delaidelli, A., Bhanshali, F., An, Y.A., Ferreira, N., Smits, L.M., Negri, G.L., Schwamborn, J.C., Jensen, P.H., et al. (2018). Activity of translation regulator eukaryotic elongation factor-2 kinase is increased in Parkinson disease brain and its inhibition reduces alpha synuclein toxicity. Acta Neuropathol Commun 6, 54. 10.1186/s40478-018-0554-9.

26. Franklin, K.B.J., and Paxinos, G. (2013). Paxinos and Franklin’s The mouse brain in stereotaxic coordinates, Fourth edition. Edition (Academic Press, an imprint of Elsevier).

27. Luo, W., and Brouwer, C. (2013). Pathview: an R/Bioconductor package for pathway-based data integration and visualization. Bioinformatics 29, 1830–1831. 10.1093/bioinformatics/btt285.

28. Watson, C., Paxinos, G., and Puelles, L. (2012). The mouse nervous system, 1st Edition (Elsevier Academic Press).

29. Bankhead, P., Loughrey, M.B., Fernandez, J.A., Dombrowski, Y., McArt, D.G., Dunne, P.D., McQuaid, S., Gray, R.T., Murray, L.J., Coleman, H.G., et al. (2017). QuPath: Open source software for digital pathology image analysis. Sci Rep 7, 16878. 10.1038/s41598-017-17204-5.

30. Anderson, J.P., Walker, D.E., Goldstein, J.M., de Laat, R., Banducci, K., Caccavello, R.J., Barbour, R., Huang, J., Kling, K., Lee, M., et al. (2006). Phosphorylation of Ser-129 is the dominant pathological modification of alpha-synuclein in familial and sporadic Lewy body disease. J Biol Chem 281, 29739–29752. 10.1074/jbc.M600933200.

31. Goedert, M., Jakes, R., and Spillantini, M.G. (2017). The Synucleinopathies: Twenty Years On. J Parkinsons Dis 7, S51–S69. 10.3233/JPD-179005.

32. Spillantini, M.G., Schmidt, M.L., Lee, V.M., Trojanowski, J.Q., Jakes, R., and Goedert, M. (1997). Alpha-synuclein in Lewy bodies. Nature 388, 839–840. 10.1038/42166.

33. Sjostedt, E., Zhong, W., Fagerberg, L., Karlsson, M., Mitsios, N., Adori, C., Oksvold, P., Edfors, F., Limiszewska, A., Hikmet, F., et al. (2020). An atlas of the protein-coding genes in the human, pig, and mouse brain. Science 367. 10.1126/science.aay5947.

34. Maniatis, S., Aijo, T., Vickovic, S., Braine, C., Kang, K., Mollbrink, A., Fagegaltier, D., Andrusivova, Z., Saarenpaa, S., Saiz-Castro, G., et al. (2019). Spatiotemporal dynamics of molecular pathology in amyotrophic lateral sclerosis. Science 364, 89–93. 10.1126/science.aav9776.

35. Gomes Moreira, D., and Jan, A. (2023). A beginner’s guide into curated analyses of open access datasets for biomarker discovery in neurodegeneration. npj Scientific Data 10, 432. 10.1038/s41597-023-02338-1.

36. Lesnick, T.G., Papapetropoulos, S., Mash, D.C., Ffrench-Mullen, J., Shehadeh, L., de Andrade, M., Henley, J.R., Rocca, W.A., Ahlskog, J.E., and Maraganore, D.M. (2007). A genomic pathway approach to a complex disease: axon guidance and Parkinson disease. PLoS Genet 3, e98. 10.1371/journal.pgen.0030098.

37. Corradini, B.R., Iamashita, P., Tampellini, E., Farfel, J.M., Grinberg, L.T., and Moreira-Filho, C.A. (2014). Complex network-driven view of genomic mechanisms underlying Parkinson’s disease: analyses in dorsal motor vagal nucleus, locus coeruleus, and substantia nigra. Biomed Res Int 2014, 543673. 10.1155/2014/543673.

38. Zheng, B., Liao, Z., Locascio, J.J., Lesniak, K.A., Roderick, S.S., Watt, M.L., Eklund, A.C., Zhang-James, Y., Kim, P.D., Hauser, M.A., et al. (2010). PGC-1alpha, a potential therapeutic target for early intervention in Parkinson’s disease. Sci Transl Med 2, 52ra73. 10.1126/scitranslmed.3001059.

39. Durrenberger, P.F., Fernando, F.S., Kashefi, S.N., Bonnert, T.P., Seilhean, D., Nait-Oumesmar, B., Schmitt, A., Gebicke-Haerter, P.J., Falkai, P., Grunblatt, E., et al. (2015). Common mechanisms in neurodegeneration and neuroinflammation: a BrainNet Europe gene expression microarray study. J Neural Transm (Vienna) 122, 1055–1068. 10.1007/s00702-014-1293-0.

40. Kel, A.E., Stegmaier, P., Valeev, T., Koschmann, J., Poroikov, V., Kel-Margoulis, O.V., and Wingender, E. (2016). Multi-omics “upstream analysis” of regulatory genomic regions helps identifying targets against methotrexate resistance of colon cancer. EuPA Open Proteom 13, 1–13. 10.1016/j.euprot.2016.09.002.

41. Nunes, I., Tovmasian, L.T., Silva, R.M., Burke, R.E., and Goff, S.P. (2003). Pitx3 is required for development of substantia nigra dopaminergic neurons. Proc Natl Acad Sci U S A 100, 4245–4250. 10.1073/pnas.0230529100.

42. Barco, A., and Kandel, E.R. (2005). The Role of CREB and CBP in Brain Function. In Transcription Factors in the Nervous System, pp. 206–241. 10.1002/3527608036.ch11.

43. Calabresi, P., Picconi, B., Tozzi, A., Ghiglieri, V., and Di Filippo, M. (2014). Direct and indirect pathways of basal ganglia: a critical reappraisal. Nat Neurosci 17, 1022–1030. 10.1038/nn.3743.

44. Lee, Y., Lee, J., Kwon, I., Nakajima, Y., Ohmiya, Y., Son, G.H., Lee, K.H., and Kim, K. (2010). Coactivation of the CLOCK-BMAL1 complex by CBP mediates resetting of the circadian clock. J Cell Sci 123, 3547–3557. 10.1242/jcs.070300.

45. Rouaux, C., Loeffler, J.P., and Boutillier, A.L. (2004). Targeting CREB-binding protein (CBP) loss of function as a therapeutic strategy in neurological disorders. Biochem Pharmacol 68, 1157–1164. 10.1016/j.bcp.2004.05.035.

46. Ettcheto, M., Abad, S., Petrov, D., Pedros, I., Busquets, O., Sanchez-Lopez, E., Casadesus, G., Beas-Zarate, C., Carro, E., Auladell, C., et al. (2018). Early Preclinical Changes in Hippocampal CREB-Binding Protein Expression in a Mouse Model of Familial Alzheimer’s Disease. Mol Neurobiol 55, 4885–4895. 10.1007/s12035-017-0690-4.

47. Julian, L., and Olson, M.F. (2014). Rho-associated coiled-coil containing kinases (ROCK): structure, regulation, and functions. Small GTPases 5, e29846. 10.4161/sgtp.29846.

48. Weber, A.J., and Herskowitz, J.H. (2021). Perspectives on ROCK2 as a Therapeutic Target for Alzheimer’s Disease. Front Cell Neurosci 15, 636017. 10.3389/fncel.2021.636017.

49. Gentry, E.G., Henderson, B.W., Arrant, A.E., Gearing, M., Feng, Y., Riddle, N.C., and Herskowitz, J.H. (2016). Rho Kinase Inhibition as a Therapeutic for Progressive Supranuclear Palsy and Corticobasal Degeneration. J Neurosci 36, 1316–1323. 10.1523/JNEUROSCI.2336-15.2016.

50. Saal, K.A., Galter, D., Roeber, S., Bahr, M., Tonges, L., and Lingor, P. (2017). Altered Expression of Growth Associated Protein-43 and Rho Kinase in Human Patients with Parkinson’s Disease. Brain Pathol 27, 13–25. 10.1111/bpa.12346.

51. Konnova, E.A., and Swanberg, M. (2018). Animal Models of Parkinson’s Disease. In Parkinson’s Disease: Pathogenesis and Clinical Aspects, T.B. Stoker, and J.C. Greenland, eds. 10.15586/codonpublications.parkinsonsdisease.2018.ch5.

52. McFarthing, K., Buff, S., Rafaloff, G., Pitzer, K., Fiske, B., Navangul, A., Beissert, K., Pilcicka, A., Fuest, R., Wyse, R.K., and Stott, S.R.W. (2024). Parkinson’s Disease Drug Therapies in the Clinical Trial Pipeline: 2024 Update. J Parkinsons Dis 14, 899–912. 10.3233/JPD-240272.

53. Uchihara, T., and Giasson, B.I. (2016). Propagation of alpha-synuclein pathology: hypotheses, discoveries, and yet unresolved questions from experimental and human brain studies. Acta Neuropathol 131, 49–73. 10.1007/s00401-015-1485-1 10.1007/s00401-015-1485-1 [pii].

54. Goedert, M., Masuda-Suzukake, M., and Falcon, B. (2017). Like prions: the propagation of aggregated tau and alpha-synuclein in neurodegeneration. Brain 140, 266–278. 10.1093/brain/aww230.

55. Hawkes, C.H., Del Tredici, K., and Braak, H. (2007). Parkinson’s disease: a dual-hit hypothesis. Neuropathol Appl Neurobiol 33, 599–614. 10.1111/j.1365-2990.2007.00874.x.

56. Surmeier, D.J., Obeso, J.A., and Halliday, G.M. (2017). Selective neuronal vulnerability in Parkinson disease. Nat Rev Neurosci 18, 101–113. 10.1038/nrn.2016.178.

57. Kamath, T., Abdulraouf, A., Burris, S.J., Langlieb, J., Gazestani, V., Nadaf, N.M., Balderrama, K., Vanderburg, C., and Macosko, E.Z. (2022). Single-cell genomic profiling of human dopamine neurons identifies a population that selectively degenerates in Parkinson’s disease. Nat Neurosci 25, 588–595. 10.1038/s41593-022-01061-1.

58. Henrich, M.T., Oertel, W.H., Surmeier, D.J., and Geibl, F.F. (2023). Mitochondrial dysfunction in Parkinson’s disease - a key disease hallmark with therapeutic potential. Mol Neurodegener 18, 83. 10.1186/s13024-023-00676-7.

59. Schapira, A.H. (2008). Mitochondria in the aetiology and pathogenesis of Parkinson’s disease. Lancet Neurol 7, 97–109. S1474-4422(07)70327-7 [pii] 10.1016/S1474-4422(07)70327-7.

60. Koprich, J.B., Johnston, T.H., Reyes, M.G., Sun, X., and Brotchie, J.M. (2010). Expression of human A53T alpha-synuclein in the rat substantia nigra using a novel AAV1/2 vector produces a rapidly evolving pathology with protein aggregation, dystrophic neurite architecture and nigrostriatal degeneration with potential to model the pathology of Parkinson’s disease. Mol Neurodegener 5, 43. 10.1186/1750-1326-5-43.

61. Cappelletti, C., Henriksen, S.P., Geut, H., Rozemuller, A.J.M., van de Berg, W.D.J., Pihlstrom, L., and Toft, M. (2023). Transcriptomic profiling of Parkinson’s disease brains reveals disease stage specific gene expression changes. Acta Neuropathol 146, 227–244. 10.1007/s00401-023-02597-7.

62. Caldi Gomes, L., Galhoz, A., Jain, G., Roser, A.E., Maass, F., Carboni, E., Barski, E., Lenz, C., Lohmann, K., Klein, C., et al. (2022). Multi-omic landscaping of human midbrains identifies disease-relevant molecular targets and pathways in advanced-stage Parkinson’s disease. Clin Transl Med 12, e692. 10.1002/ctm2.692.

63. Zhang, Y., James, M., Middleton, F.A., and Davis, R.L. (2005). Transcriptional analysis of multiple brain regions in Parkinson’s disease supports the involvement of specific protein processing, energy metabolism, and signaling pathways, and suggests novel disease mechanisms. Am J Med Genet B Neuropsychiatr Genet 137B, 5–16. 10.1002/ajmg.b.30195.

64. Fiorini, M.R., Dilliott, A.A., Thomas, R.A., and Farhan, S.M.K. (2024). Transcriptomics of Human Brain Tissue in Parkinson’s Disease: a Comparison of Bulk and Single-cell RNA Sequencing. Mol Neurobiol. 10.1007/s12035-024-04124-5.

65. Bolinger, A.A., Frazier, A., La, J.H., Allen, J.A., and Zhou, J. (2023). Orphan G Protein-Coupled Receptor GPR37 as an Emerging Therapeutic Target. ACS Chem Neurosci 14, 3318–3334. 10.1021/acschemneuro.3c00479.

66. Naia, L., Shimozawa, M., Bereczki, E., Li, X., Liu, J., Jiang, R., Giraud, R., Leal, N.S., Pinho, C.M., Berger, E., et al. (2023). Mitochondrial hypermetabolism precedes impaired autophagy and synaptic disorganization in App knock-in Alzheimer mouse models. Mol Psychiatry 28, 3966–3981. 10.1038/s41380-023-02289-4.

67. Delaidelli, A., Jan, A., Herms, J., and Sorensen, P.H. (2019). Translational control in brain pathologies: biological significance and therapeutic opportunities. Acta Neuropathologica 137, 535–555. 10.1007/s00401-019-01971-8.

68. Garcia-Esparcia, P., Hernandez-Ortega, K., Koneti, A., Gil, L., Delgado-Morales, R., Castano, E., Carmona, M., and Ferrer, I. (2015). Altered machinery of protein synthesis is region- and stage-dependent and is associated with alpha-synuclein oligomers in Parkinson’s disease. Acta Neuropathol Commun 3, 76. 10.1186/s40478-015-0257-4.

69. Salminen, A., and Kaarniranta, K. (2012). AMP-activated protein kinase (AMPK) controls the aging process via an integrated signaling network. Ageing Res Rev 11, 230–241. 10.1016/j.arr.2011.12.005.

70. Crino, P.B. (2016). The mTOR signalling cascade: paving new roads to cure neurological disease. Nat Rev Neurol 12, 379–392. 10.1038/nrneurol.2016.81.

71. Sossin, W.S., and Costa-Mattioli, M. (2019). Translational Control in the Brain in Health and Disease. Cold Spring Harb Perspect Biol 11. 10.1101/cshperspect.a032912.

72. Cappellano, G., Vecchio, D., Magistrelli, L., Clemente, N., Raineri, D., Barbero Mazzucca, C., Virgilio, E., Dianzani, U., Chiocchetti, A., and Comi, C. (2021). The Yin-Yang of osteopontin in nervous system diseases: damage versus repair. Neural Regen Res 16, 1131–1137. 10.4103/1673-5374.300328.

73. Koch, J.C., Tatenhorst, L., Roser, A.E., Saal, K.A., Tonges, L., and Lingor, P. (2018). ROCK inhibition in models of neurodegeneration and its potential for clinical translation. Pharmacol Ther 189, 1–21. 10.1016/j.pharmthera.2018.03.008.

74. Schapira, A.H.V., Chaudhuri, K.R., and Jenner, P. (2017). Non-motor features of Parkinson disease. Nat Rev Neurosci 18, 435–450. 10.1038/nrn.2017.62.

75. Postuma, R.B., Gagnon, J.F., Bertrand, J.A., Genier Marchand, D., and Montplaisir, J.Y. (2015). Parkinson risk in idiopathic REM sleep behavior disorder: preparing for neuroprotective trials. Neurology 84, 1104–1113. 10.1212/WNL.0000000000001364.

76. Videnovic, A., and Golombek, D. (2013). Circadian and sleep disorders in Parkinson’s disease. Exp Neurol 243, 45–56. 10.1016/j.expneurol.2012.08.018.

77. Chen, Y.C., Wang, W.S., Lewis, S.J.G., and Wu, S.L. (2024). Fighting Against the Clock: Circadian Disruption and Parkinson’s Disease. J Mov Disord 17, 1–14. 10.14802/jmd.23216.

78. Butkovich, L.M., Houser, M.C., Chalermpalanupap, T., Porter-Stransky, K.A., Iannitelli, A.F., Boles, J.S., Lloyd, G.M., Coomes, A.S., Eidson, L.N., De Sousa Rodrigues, M.E., et al. (2020). Transgenic Mice Expressing Human alpha-Synuclein in Noradrenergic Neurons Develop Locus Ceruleus Pathology and Nonmotor Features of Parkinson’s Disease. J Neurosci 40, 7559–7576. 10.1523/JNEUROSCI.1468-19.2020.

79. Kudo, T., Loh, D.H., Truong, D., Wu, Y., and Colwell, C.S. (2011). Circadian dysfunction in a mouse model of Parkinson’s disease. Exp Neurol 232, 66–75. 10.1016/j.expneurol.2011.08.003.

80. Rothman, S.M., Griffioen, K.J., Vranis, N., Ladenheim, B., Cong, W.N., Cadet, J.L., Haran, J., Martin, B., and Mattson, M.P. (2013). Neuronal expression of familial Parkinson’s disease A53T alpha-synuclein causes early motor impairment, reduced anxiety and potential sleep disturbances in mice. J Parkinsons Dis 3, 215–229. 10.3233/JPD-120130.

81. Theologidis, V., Ferreira, S.A., Jensen, N.M., Moreira, D.G., Ahlgreen, O.A., Hansen, M.W., Rosenberg, E.D., Richner, M., Faress, I., Gram, H., et al. (2024). Bradykinesia and postural instability in a model of prodromal Synucleinopathy with α-Synuclein aggregation in the gigantocellular nuclei. bioRxiv, 2024.2009.2005.610956. 10.1101/2024.09.05.610956.

82. Chen, W.T., Lu, A., Craessaerts, K., Pavie, B., Sala Frigerio, C., Corthout, N., Qian, X., Lalakova, J., Kuhnemund, M., Voytyuk, I., et al. (2020). Spatial Transcriptomics and In Situ Sequencing to Study Alzheimer’s Disease. Cell 182, 976–991 e919. 10.1016/j.cell.2020.06.038.

83. Goralski, T.M., Meyerdirk, L., Breton, L., Brasseur, L., Kurgat, K., DeWeerd, D., Turner, L., Becker, K., Adams, M., Newhouse, D.J., and Henderson, M.X. (2024). Spatial transcriptomics reveals molecular dysfunction associated with cortical Lewy pathology. Nat Commun 15, 2642. 10.1038/s41467-024-47027-8.

84. Swarup, V., Hinz, F.I., Rexach, J.E., Noguchi, K.I., Toyoshiba, H., Oda, A., Hirai, K., Sarkar, A., Seyfried, N.T., Cheng, C., et al. (2019). Identification of evolutionarily conserved gene networks mediating neurodegenerative dementia. Nat Med 25, 152–164. 10.1038/s41591-018-0223-3.

85. van den Hurk, M., Lau, S., Marchetto, M.C., Mertens, J., Stern, S., Corti, O., Brice, A., Winner, B., Winkler, J., Gage, F.H., and Bardy, C. (2022). Druggable transcriptomic pathways revealed in Parkinson’s patient-derived midbrain neurons. NPJ Parkinsons Dis 8, 134. 10.1038/s41531-022-00400-0.

