## Supplementary material for "Molecular signatures of altered energy metabolism and circadian rhythm perturbations in a model of extra-nigral Synucleinopathy": Supplemenatry Information Lin et al.

#### SUPPLEMENTARY INFORMATION

- Table S1: Demographics, human cases used for IHC
- Supplementary Figures Legends
- Supplementary Figures
- User Guide for the online ST database

| Table S1. Demographics of the cases examined for IHC studies | | | | | |
|---|---|---|---|---|---|
| **Case ID** | **Age** | **Gender** | **Clinical Diagnosis** | **LBD** | **Remarks** |
| PD-1 | 71 | Male | PD | limbic | TDP-43 pathology in limbic structures |
| PD-2 | 69 | Male | PD | diffuse | |
| PD-3 | 87 | Male | PD + dementia | diffuse | |
| PD-4 | 74 | Male | PD + dementia | diffuse | |
| PD-5 | 67 | Male | PD | diffuse | acute anoxic-ischemia |
| PD-6 | 74 | Male | PD | diffuse | |
| C-1 | 78 | Male | CVD | none | CVD |
| C-2 | 93 | Female | CVD | none | CVD |
| C-3 | 76 | Female | CVD | none | |
| C-4 | 50 | Male | CVD | none | |
| C-5 | 76 | Male | CVD | none | |
| C-6 | 71 | Female | Lymphoma | none | |

**Abbreviations:** C= Control; CVD: chronic cerebrovascular disease; LBD = Lewy body disease/Pathology; PD = Parkinson disease
LBD stage: brainstem (brainstem only), limbic (brainstem + limbic), diffuse (brainstem + limbic + neocortex)

#### SUPPLEMENTARY FIGURE LEGENDS

**S1. Schematic depictions of select KEGG metabolic pathways in M83$^{+/+}$ ST data. (A-C)** Pathway maps depicting the relative expression of transcripts in select KEGG pathways: glycolysis/gluconeogenesis (in A), oxidative phosphorylation (in B) and fatty acid metabolism (in C), in pair-wise comparisons involving Early stage (E) vs. Controls (C) and Late stage (L) vs. Early stage (E) cohorts of M83$^{+/+}$ mice - Also see Fig. 3-4. Pathways were visualized using Pathview R package (see Methods).

**S2. Schematic depictions of select KEGG pathways in M83$^{+/+}$ ST data. (A-C)** Pathway maps depicting the relative expression of transcripts in select KEGG pathways: ribosome assembly (in A), immune system/antigen presentation (in B) and actin-mediated regulation of cytoskeleton (in C), in pair-wise comparisons involving Early stage (E) vs. Controls (C) and Late stage (L) vs. Early stage (E) cohorts of M83$^{+/+}$ mice - Also see Fig. 3 and Fig. 5. Pathways were visualized using Pathview R package (see Methods).

**S3. Heatmaps depicting the relative expression of select ST transcripts in M83$^{+/+}$ brain and PD microarray datasets. (A)** Heatmaps depicting the relative expression of select 50 transcripts in ST data in pair-wise comparisons involving Early stage (E) vs. Controls (C) and Late stage (L) vs. Early stage (E) cohorts of M83$^{+/+}$ mice. **(B-C)** Heatmaps depicting the relative expression of select 50 transcripts in PD microarray datasets, with Log2 fold-change and respective p-value in a given dataset. The datasets and brain regions examined included: 1) GSE7621 (*substantia nigra-SN*; Controls, n=9; PD, n=16), 2) GSE43490 (*SN, dorsal motor nucleus of vagus- dmX and locus coeruleus-LC*; Controls, n=5-7; PD, n=8), 3) GSE20146 (*globus pallidus, interna-GPi*; Controls, n=10; PD, n=10) and 4) GSE26927 (*SN*; Controls, n=7; PD, n=12). Notice the *Crebbp/CREBBP* and *Rock2/ROCK2* expression in relation to aSyn pathology in (A) and in across the PD microarray datasets (in B).

**S4. Heatmaps depicting the relative enrichment of transcription factor (TF) regulons in relation to the stage of aSyn pathology in brains of M83$^{+/+}$ mice. (A-B)** Heatmaps depicting the relative enrichment of TF regulons in ST data in pair-wise comparisons involving Early stage (E) vs. Controls (C) and Late stage (L) vs. Early stage (E) cohorts of M83$^{+/+}$ mice. The TF regulons are grouped according to enrichment in the early stage- ES (in A) or late stage-LS of aSyn pathology. Abbreviations: Crb (cerebellum), Ch. plx (choroid plexus), Ctx (cerebral cortex), hyp. th. (hypothalamus), Mb (midbrain), Thal. (thalamus) and wh. mat. (white matter). Also see Fig. 3-5.

**S5. ST maps and IF detection of ROCK2 in brains of M83$^{+\backslash-}$ mice. (A)** Representative spatial maps depicting the relative abundance of *Rock2* in sagittal brain sections from cohorts of M83$^{+/+}$ mice. **(B)** Representative (20X) images showing ROCK2 IF in the gigantocellular nuclei (GRN, in pons), periaqueductal grey (PAG, in midbrain), deep cerebellar nuclei (DCN, in cerebellum) and mediodorsal nuclei (MD, in thalamus). The inset show 63X magnified views from the regions. Scale bar= 50 µm. **(C)** Bar graphs depicting quantification of ROCK2 IF in GRN, PAG, DCN and DM of the experimental cohorts, as indicated. Error bars depict Mean IF intensity ± SD as % of cells in total area (PBS, n=2; DPI-45, n=3 and DPI-75, n=3; See Methods). Statistics in S5C: Kruskal-Wallis ANOVA (p-values on graphs), Dunn's multiple comparisons (non-significant).

**S6. Quantitation of p-aSyn (S129) and CREBBP/CBP IHC in post-mortem human brain sections. (A-B)** Bar graphs depicting quantification of p-aSyn, S129 and CREBBP/CBP IHC in the SN (in A) and PAG (in B) of controls (C1-C6) and PD cases (PD1-PD6) examined in the study. Graphs display Mean ± SEM of Lewy bodies per mm$^2$ and Mean % ± SEM of CREBBP/CBP immunopositive cells in the region (see Methods). Notice a relatively higher detection of CREBBP in PD-3 and PD-4, cases with diffuse LB + dementia (Table S1). Also see Fig. 7A-F.

### SELECT KEGG PATHWAYS

**Early stage aSyn pathology** | **Late stage aSyn pathology**

#### A) Glycolylsis/Gluconeogenesis (mmu00010)

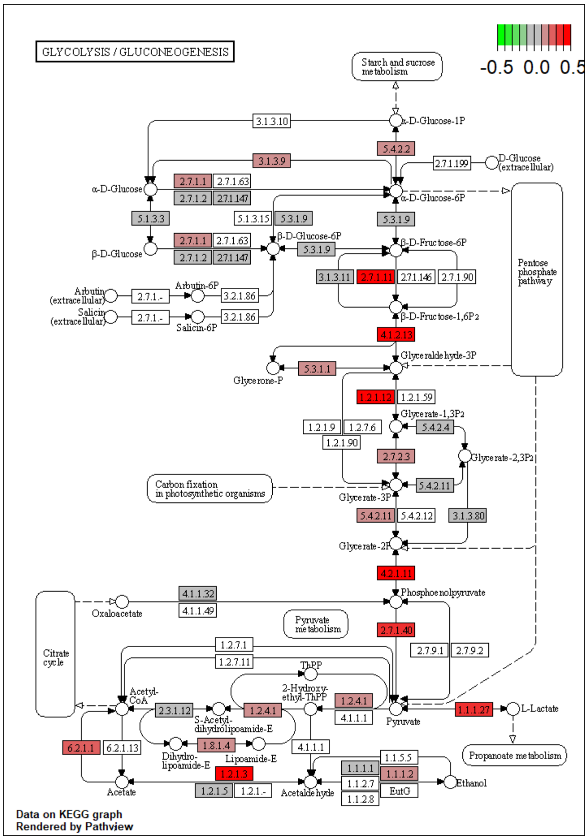


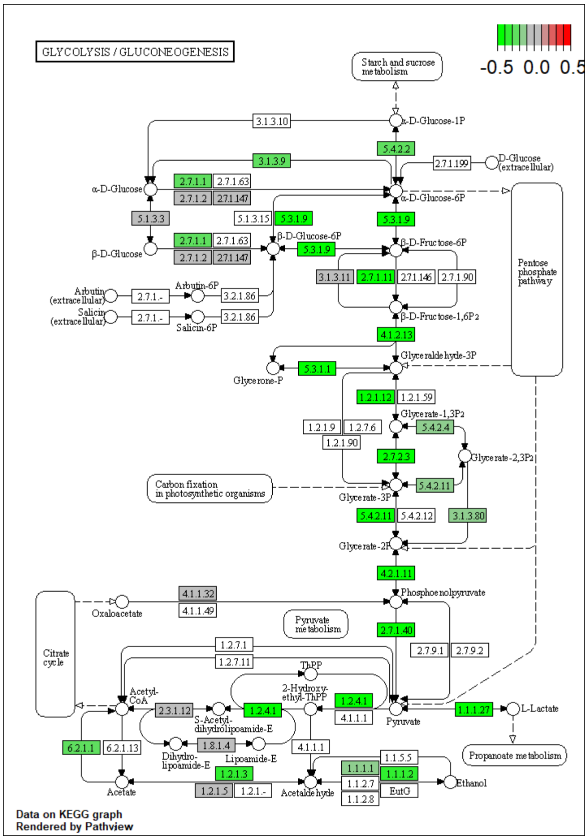


#### B) Oxidative Phosphorylation (mmu00190)

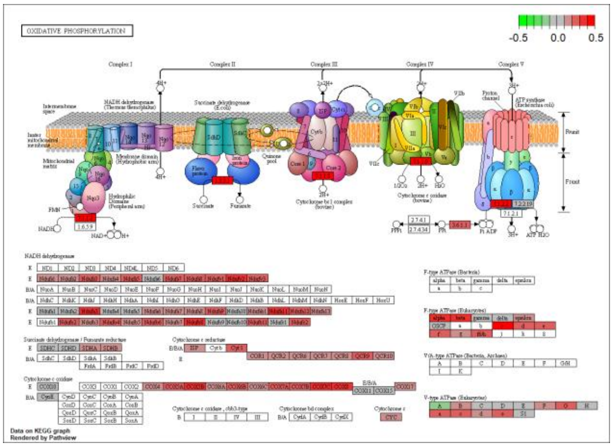


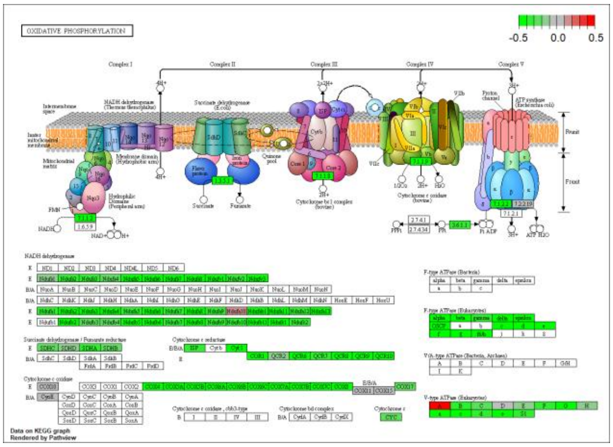


#### C) PPARγ (mmu03320)

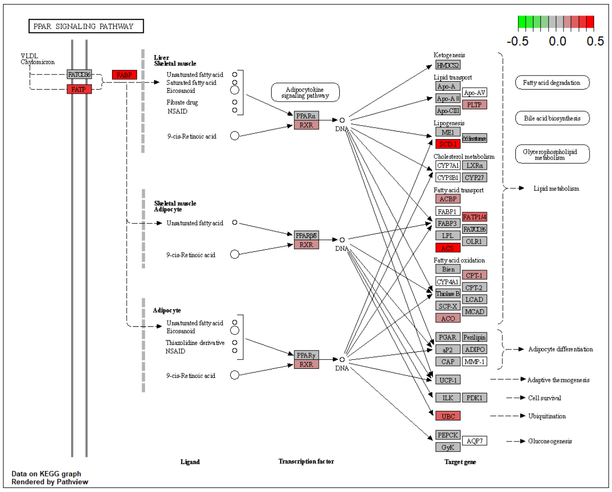


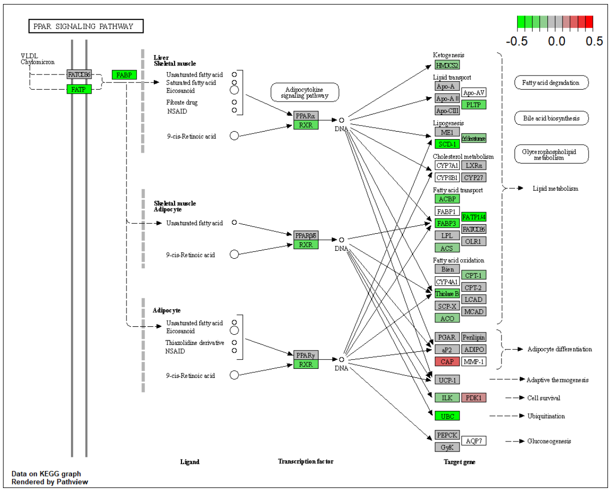

### SELECT KEGG PATHWAYS

#### Early stage aSyn pathology | Late stage aSyn pathology

#### A) Ribosome (mmu03010)

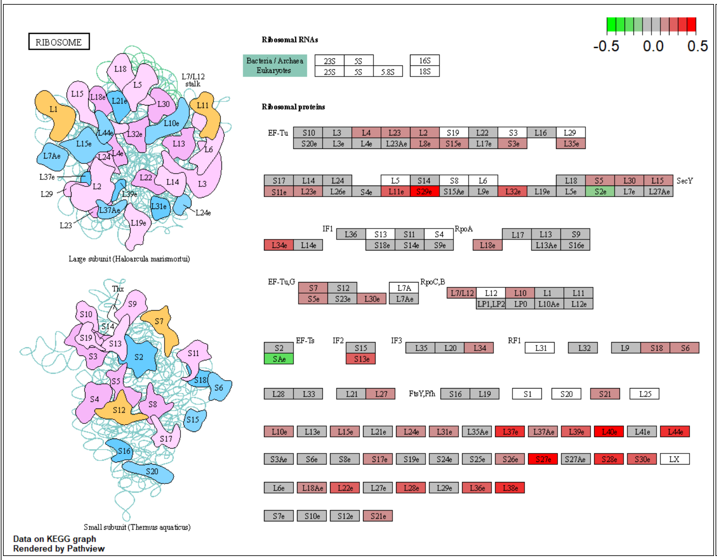


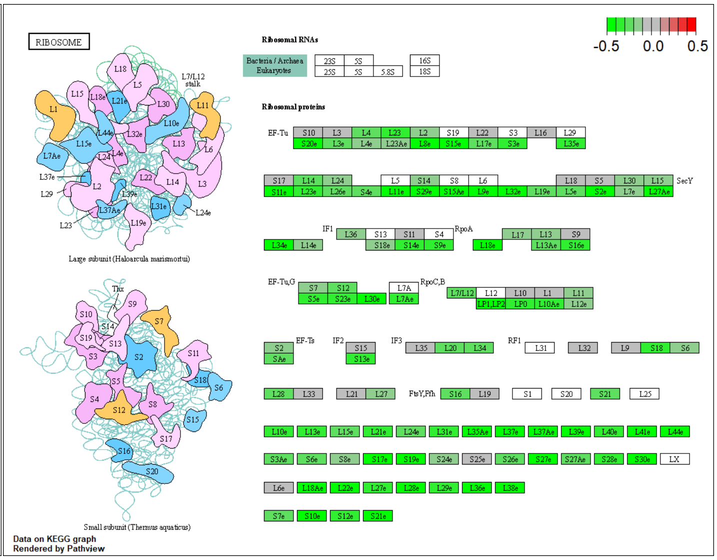


#### B) Antigen processing and presentation (mmu04612)

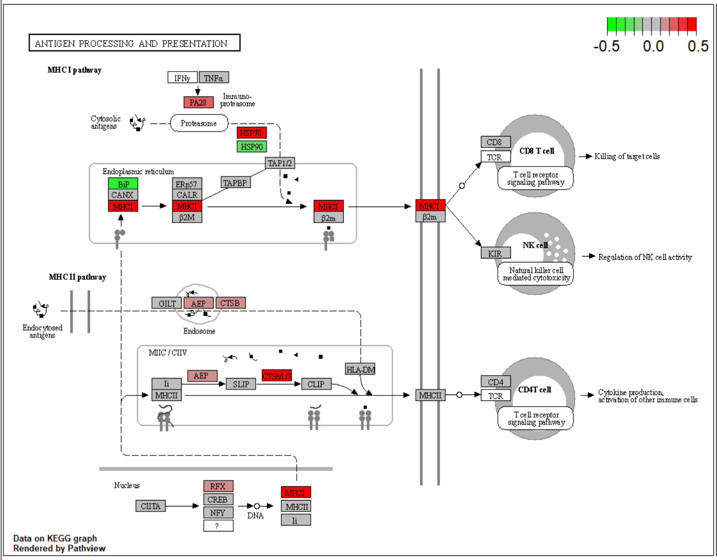


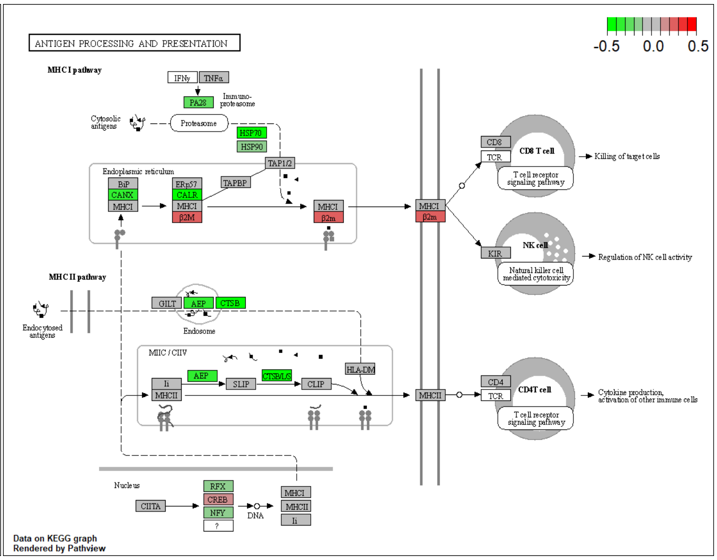


#### C) Regulation of actin cytoskeleton (mmu04810)

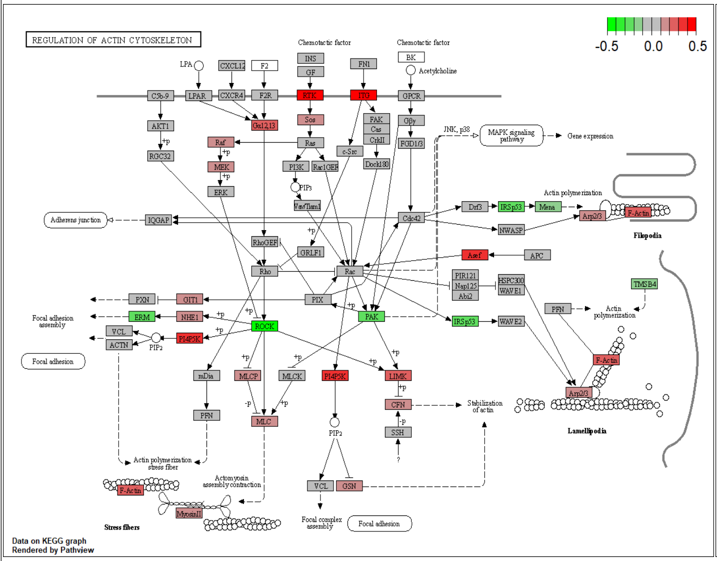


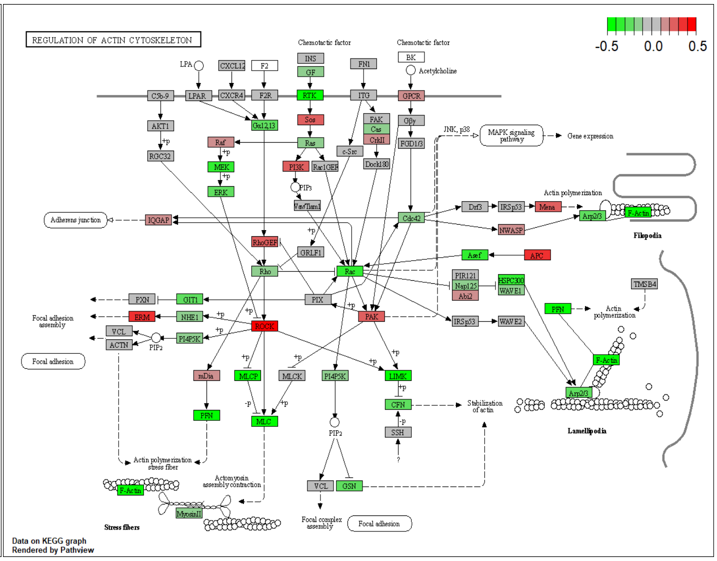

S3

A) ST profiles (aSyn M83 mouse model)

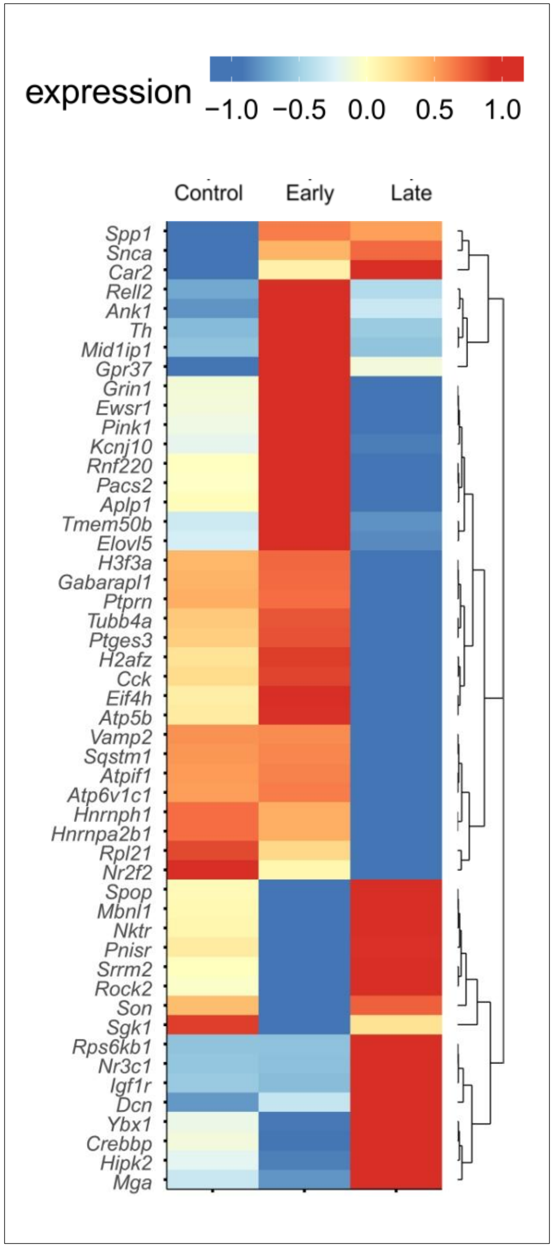


B) PD Microarray Expression Profiles

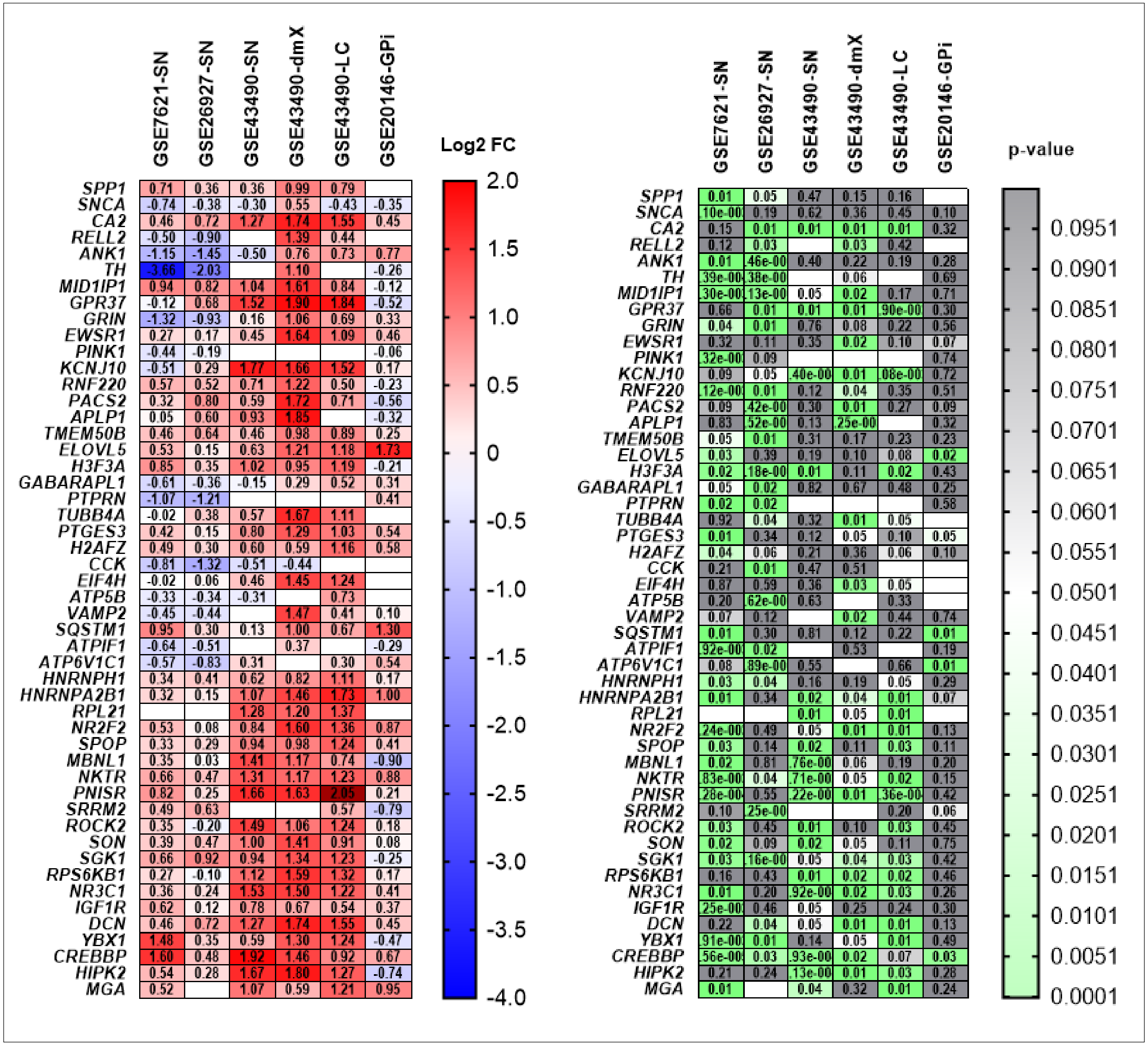

S4

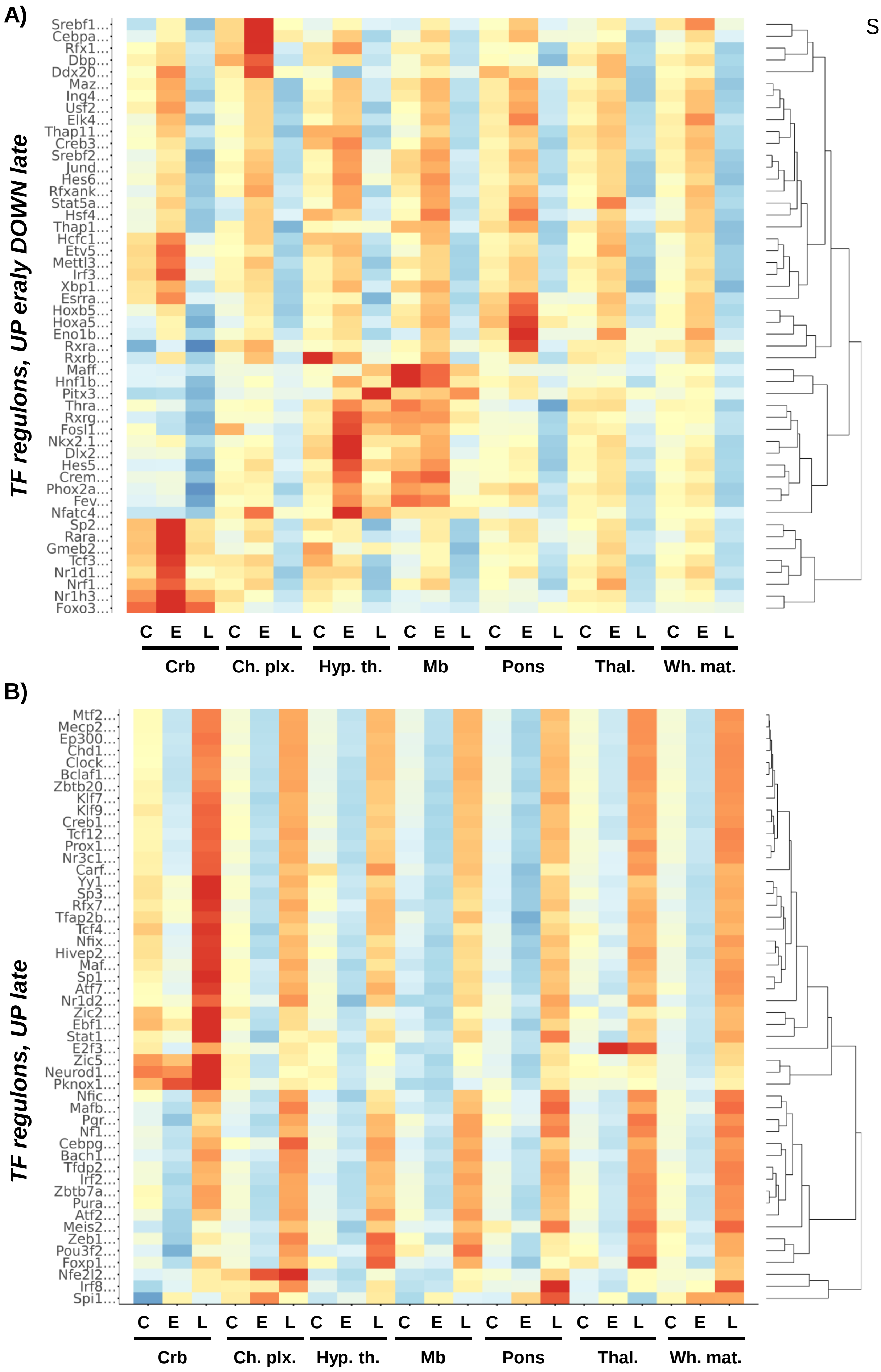
A)
TF regulons, UP eraly DOWN late
Srebf1...
Cebpa...
Rfx1...
Dbp...
Ddx20...
Maz...
Ing4...
Usf2...
Elk4...
Thap11...
Creb3...
Srebf2...
Jund...
Hes6...
Rfxank...
Stat5a...
Hsf4...
Thap1...
Hcfc1...
Etv5...
Mettl3...
Irf3...
Xbp1...
Esrra...
Hoxb5...
Hoxa5...
Eno1b...
Rxra...
Rxrb...
Maff...
Hnf1b...
Pitx3...
Thra...
Rxrg...
Fosl1...
Nkx2.1...
Dlx2...
Hes5...
Crem...
Phox2a...
Fev...
Nfatc4...
Sp2...
Rara...
Gmeb2...
Tcf3...
Nr1d1...
Nrf1...
Nr1h3...
Foxo3...
C E L C E L C E L C E L C E L C E L C E L
Crb
Ch. plx.
Hyp. th.
Mb
Pons
Thal.
Wh. mat.
B)
TF regulons, UP late
Mtf2...
Mecp2...
Ep300...
Chd1...
Clock...
Bclaf1...
Zbtb20...
Klf7...
Klf9...
Creb1...
Tcf12...
Prox1...
Nr3c1...
Carf...
Yy1...
Sp3...
Rfx7...
Tfap2b...
Tcf4...
Nfix...
Hivep2...
Maf...
Sp1...
Atf7...
Nr1d2...
Zic2...
Ebf1...
Stat1...
E2f3...
Zic5...
Neurod1...
Pknox1...
Nfic...
Mafb...
Pgr...
Nf1...
Cebpg...
Bach1...
Tfdp2...
Irf2...
Zbtb7a...
Pura...
Atf2...
Meis2...
Zeb1...
Pou3f2...
Foxp1...
Nfe2l2...
Irf8...
Spi1...
C E L C E L C E L C E L C E L C E L C E L
Crb
Ch. plx.
Hyp. th.
Mb
Pons
Thal.
Wh. mat.

S5

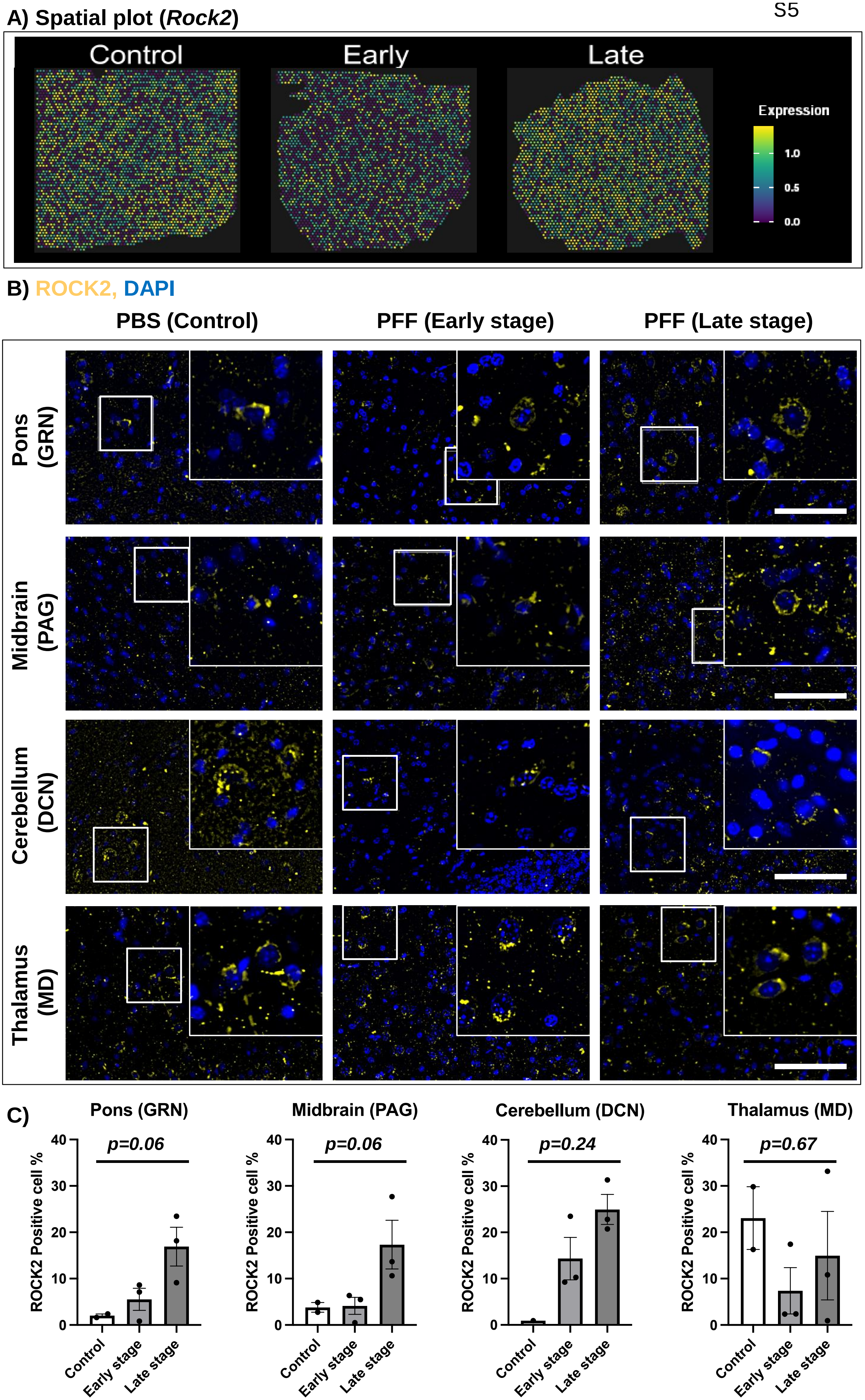

A) Spatial plot (Rock2)
Control
Early
Late
Expression
1.0
0.5
0.0
B) ROCK2, DAPI
PBS (Control)
PFF (Early stage)
PFF (Late stage)
Pons (GRN)
Midbrain (PAG)
Cerebellum (DCN)
Thalamus (MD)
C)
Pons (GRN)
Midbrain (PAG)
Cerebellum (DCN)
Thalamus (MD)
ROCK2 Positive cell %
p=0.06
p=0.06
p=0.24
p=0.67
0
10
20
30
40
Control
Early stage
Late stage

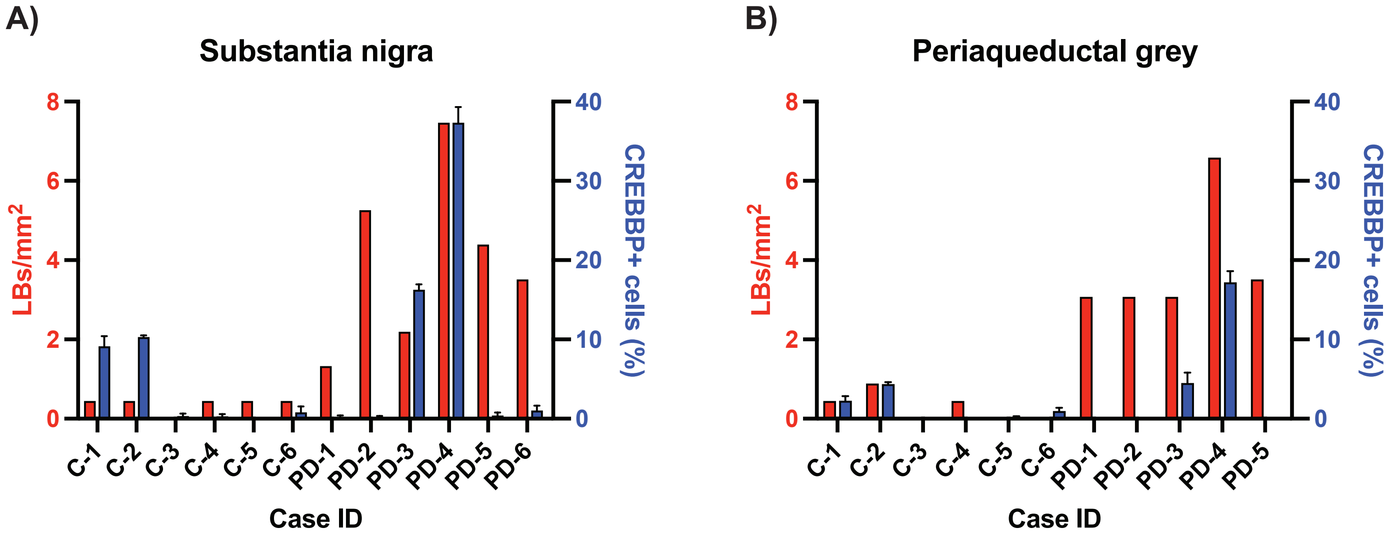
A)
Substantia nigra
LBs/mm2
CREBBP+ cells (%)
8
6
4
2
0
40
30
20
10
0
C-1
C-2
C-3
C-4
C-5
C-6
PD-1
PD-2
PD-3
PD-4
PD-5
PD-6
Case ID
B)
Periaqueductal grey
LBs/mm2
CREBBP+ cells (%)
8
6
4
2
0
40
30
20
10
0
C-1
C-2
C-3
C-4
C-5
C-6
PD-1
PD-2
PD-3
PD-4
PD-5
Case ID

#### Introduction to the Synucleinopathy M83 model Spatial Database

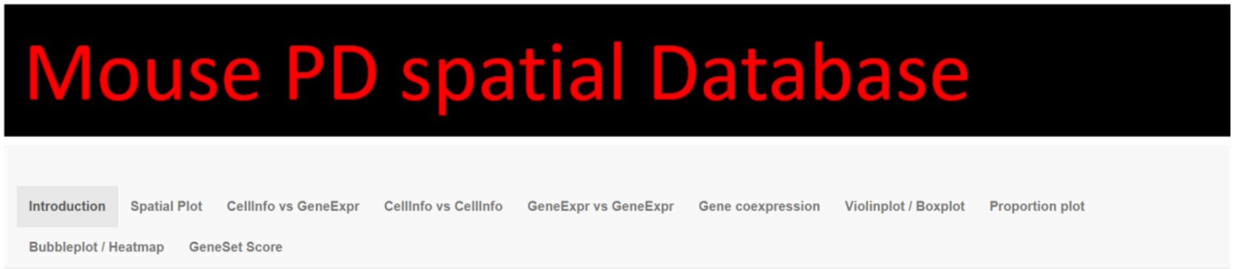


The Mouse PD spatial database is designed to help researchers and clinicians explore gene expression profiles in the context of Parkinson's disease mouse models. Here's a practical guide on how to navigate and utilize the database effectively:

**Accessing the Database**

https://dreamapp.biomed.au.dk/PD_spatial_mouse_DB/

**Navigating the Interface**: Use the tabs at the top of the page to switch between different sections of the database: Introduction, Spatial Plot, CellInfo vs GeneExpr, CellInfo vs CellInfo, GeneExpr vs GeneExpr, Gene coexpression, Violinplot/ Boxplot, Proportion plot, Bubbleplot/ Heatmap, GeneSet Score.

**Data Visualization:**

1. Spatial Plots
2. UMAPs
3. Violin and Box plots
4. Cell Proportions
5. Heatmaps and Bubble plots
6. Gene set scores (HALLMARK/KEGG)

#### 1. How to Use the Spatial Plot Tab

The Spatial Plot tab in the Mouse Parkinson's Disease (PD) spatial database allows users to explore the genes that exhibit significant changes in expression levels across different brain regions and different time points (early PD or late PD vs. sham conditions). Here's a detailed guide on how to use this feature:

##### <u>1.1. Accessing Spatial gene expression</u>

**1.1.1. Select the Spatial Plot Tab**: Click on the "Spatial Plot" tab on the top navigation bar to open the differentially expressed genes analysis interface.

##### <u>1.2 Searching for a Specific Gene</u>

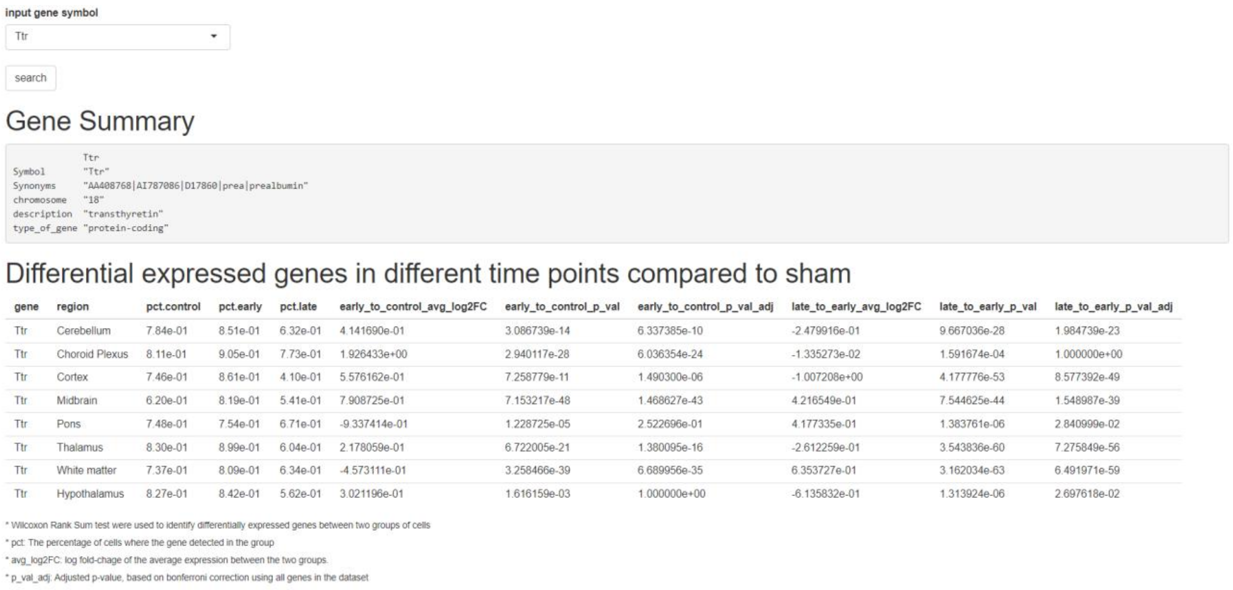
input gene symbol

Ttr

search

Gene Summary

```
            Ttr
Symbol      "Ttr"
Synonyms    "AA408768|AI787086|D17860|prea|prealbumin"
chromosome  "18"
description "transthyretin"
type_of_gene "protein-coding"
```

Differential expressed genes in different time points compared to sham

| gene | region | pct.control | pct.early | pct.late | early_to_control_avg_log2FC | early_to_control_p_val | early_to_control_p_val_adj | late_to_early_avg_log2FC | late_to_early_p_val | late_to_early_p_val_adj |
|---|---|---|---|---|---|---|---|---|---|---|
| Ttr | Cerebellum | 7.84e-01 | 8.51e-01 | 6.32e-01 | 4.141690e-01 | 3.086739e-14 | 6.337385e-10 | -2.479916e-01 | 9.667036e-28 | 1.984739e-23 |
| Ttr | Choroid Plexus | 8.11e-01 | 9.05e-01 | 7.73e-01 | 1.926433e+00 | 2.940117e-28 | 6.036354e-24 | -1.335273e-02 | 1.591674e-04 | 1.000000e+00 |
| Ttr | Cortex | 7.46e-01 | 8.61e-01 | 4.10e-01 | 5.576162e-01 | 7.258779e-11 | 1.490300e-06 | -1.007208e+00 | 4.177776e-53 | 8.577392e-49 |
| Ttr | Midbrain | 6.20e-01 | 8.19e-01 | 5.41e-01 | 7.908725e-01 | 7.153217e-48 | 1.468627e-43 | 4.216549e-01 | 7.544625e-44 | 1.548987e-39 |
| Ttr | Pons | 7.48e-01 | 7.54e-01 | 6.71e-01 | -9.337414e-01 | 1.228725e-05 | 2.522696e-01 | 4.177335e-01 | 1.383761e-06 | 2.840999e-02 |
| Ttr | Thalamus | 8.30e-01 | 8.99e-01 | 6.04e-01 | 2.178059e-01 | 6.722005e-21 | 1.380095e-16 | -2.612259e-01 | 3.543836e-60 | 7.275849e-56 |
| Ttr | White matter | 7.37e-01 | 8.09e-01 | 6.34e-01 | -4.573111e-01 | 3.258466e-39 | 6.689956e-35 | 6.353727e-01 | 3.162034e-63 | 6.491971e-59 |
| Ttr | Hypothalamus | 8.27e-01 | 8.42e-01 | 5.62e-01 | 3.021196e-01 | 1.616159e-03 | 1.000000e+00 | -6.135832e-01 | 1.313924e-06 | 2.697618e-02 |

* Wilcoxon Rank Sum test were used to identify differentially expressed genes between two groups of cells
* pct: The percentage of cells where the gene detected in the group
* avg_log2FC: log fold-chage of the average expression between the two groups.
* p_val_adj: Adjusted p-value, based on bonferroni correction using all genes in the dataset

**1.2.1. Input Gene Symbol**:

- Type the gene symbol of interest in the input box (e.g., "Ttr").
- Click the "search" button to retrieve the gene expression data.

**1.2.2. Gene Summary**:

- After searching, a summary of the gene will be displayed, including its symbol, synonyms, chromosome location, description, and type

**1.2.3. View Results**:

- The results will be displayed in a table, showing various columns such as:
  - **Gene**: The selected gene, in this example the transthyretin (Ttr) gene
  - **Region**: The specific brain region selected.
  - **Pct.control**: The percentage of cells where the gene was detected in the control group
  - **Pct.early**: The percentage of cells expressing the gene in the early PD group
  - **Pct.late**: The percentage of cells expressing the gene in the late PD group

- **avg_log2FC**: The average log2 fold change in expression in the different conditions (early_to_control and late_to_early).
- **P_val**: The p-value indicating the statistical significance of the expression change.
- **P_val_adj**: The adjusted p-value for multiple comparisons.

#### 1.3. Visualizing Spatial Gene Expression

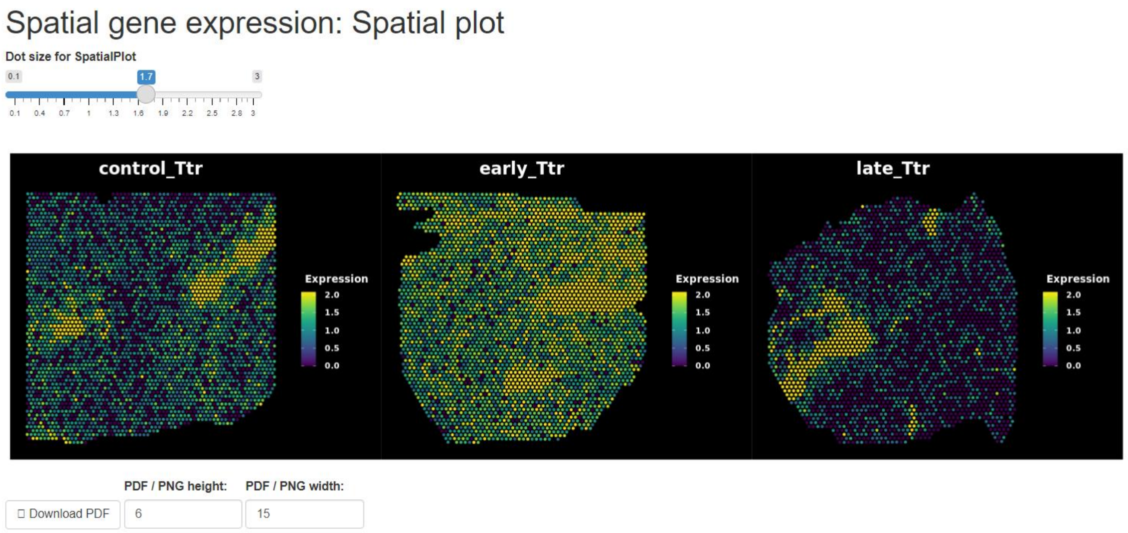


##### 1.3.1. Spatial gene expression: Spatial Plot:

- This section provides a visual representation of the gene expression in spatial context. You can see the selected gene expression in different time points and different brain regions (vs. sham), together with their corresponding spatial plots
- Adjust the dot size using the slider to change the visualization resolution.
- Download the spatial plot as a PDF or PNG file by setting the desired dimensions and clicking the download buttons.

#### 2. Analyzing Gene Expression on Reduced Dimensions

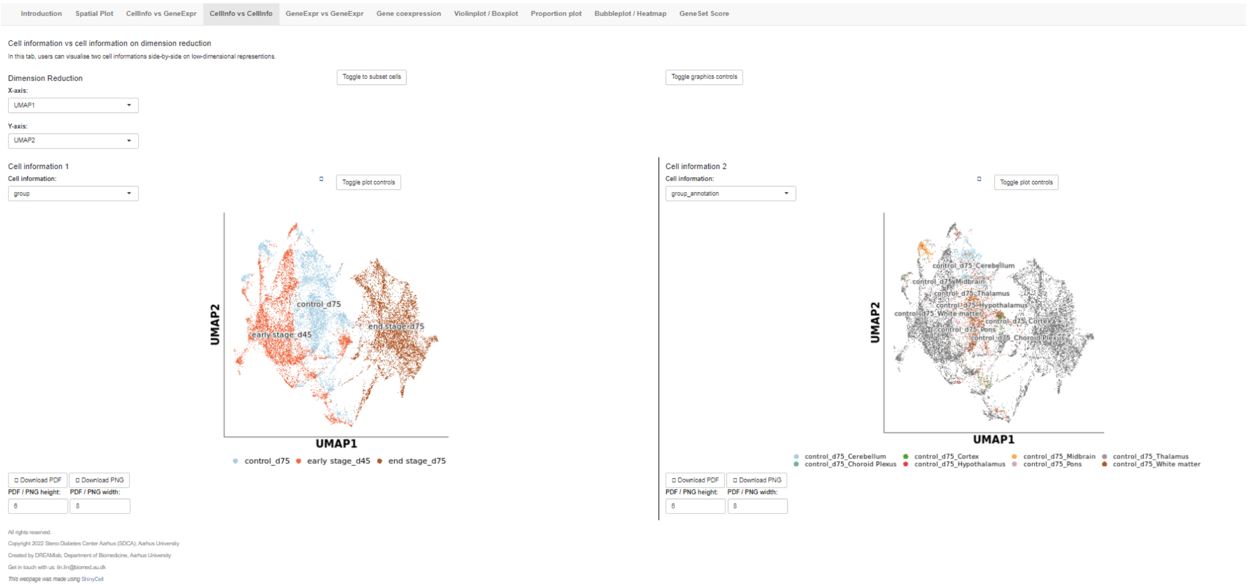


##### 2.1. Dimension Reduction Plot – CellInfo vs GeneExpr

- Use this section to visualize gene expression on reduced dimensions (e.g., UMAP);
- Select the axes for dimension reduction (e.g., UMAP1 and UMAP2);
- Choose the primary cell information you are interested;
- Click on “Toggle to subset cells” to toggle between different spot information and gene expression views. Here you can select which cells to show by selecting “Cell information to subset”;
- On the right panel select the gene you would like to visualize on the UMAP plot;
- Download the plots as PDF or PNG files.

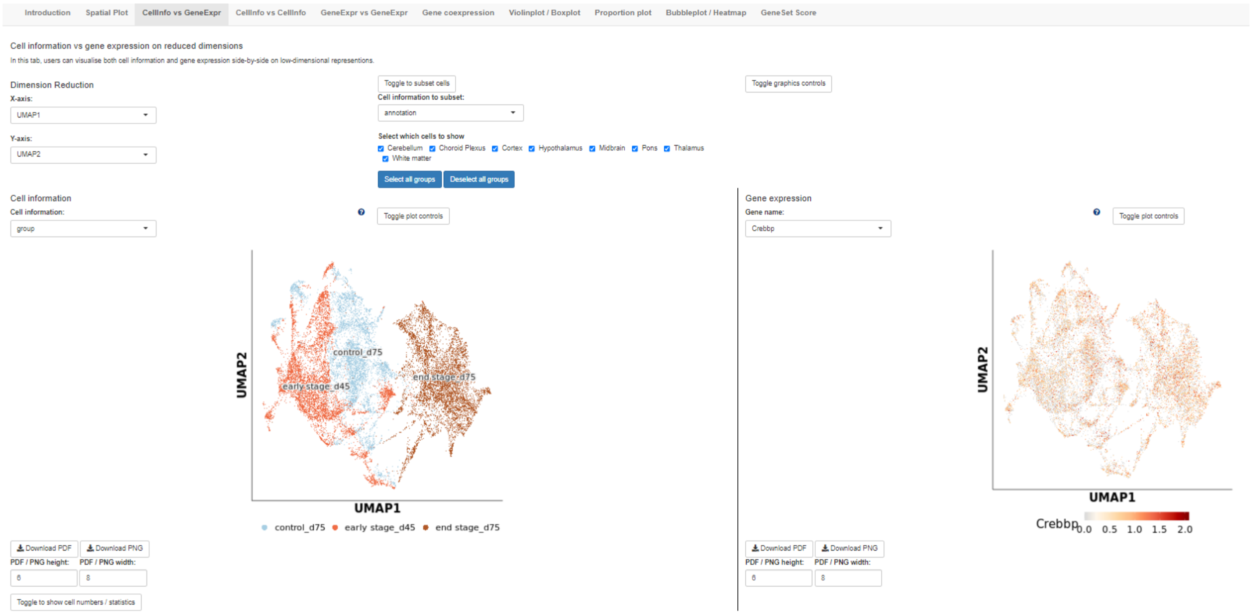


#### 2.2. Dimension Reduction Plot –CellInfo vs CellInfo

- Similarly to above, use this section to visualize cell information on reduced dimensions (e.g., UMAP);
- Select the axes for dimension reduction (e.g., UMAP1 and UMAP2);
- Toggle between different spot information and cell information views;
- Download the plots as PDF or PNG files.

#### 2.3. Co-expression of Two Genes

##### 2.3.1. Dimension Reduction Plot –GeneExpr vs GeneExpr:

- Use this section to visualize gene co-expression on reduced dimensions (e.g., UMAP);
- Select the axes for dimension reduction (e.g., UMAP1 and UMAP2);
- Toggle between different spot information and gene expression views;
- Download the plots as PDF or PNG files.

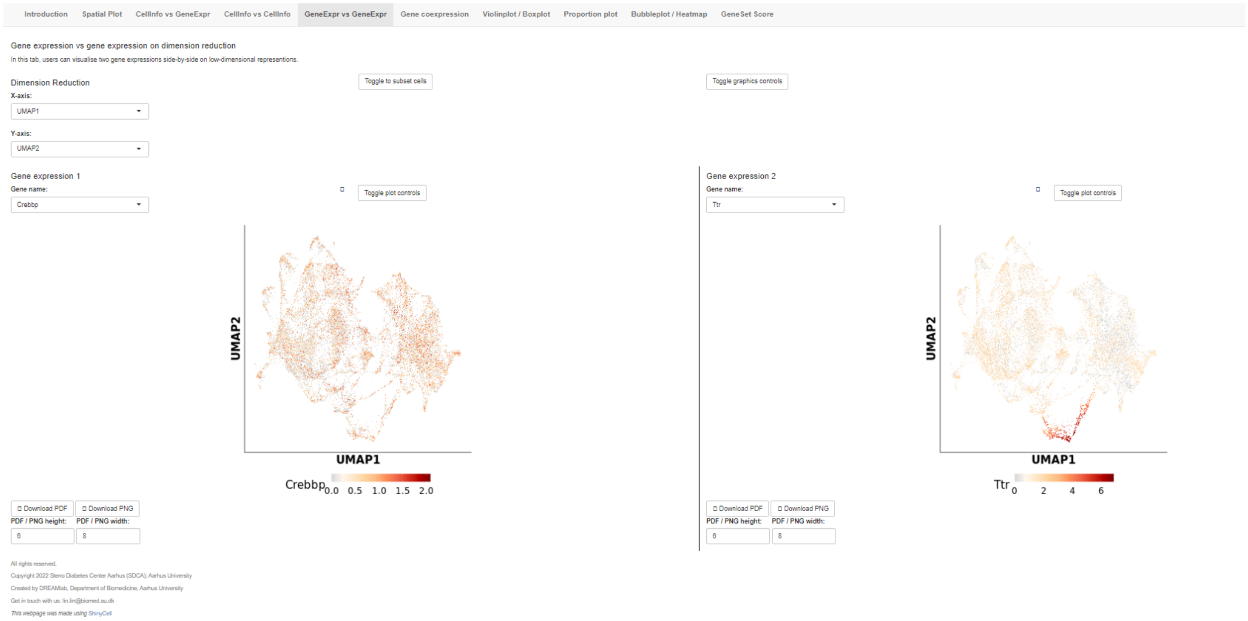


##### 2.3.2. Co-expression Plot:

- This section allows you to analyze the co-expression of two genes on reduced dimensions;
- Select the genes to be analyzed (e.g., "Crebbp" and "Ttr");
- The plot shows the spatial distribution of cells expressing both genes;
- Download the co-expression plot as a PDF or PNG file.

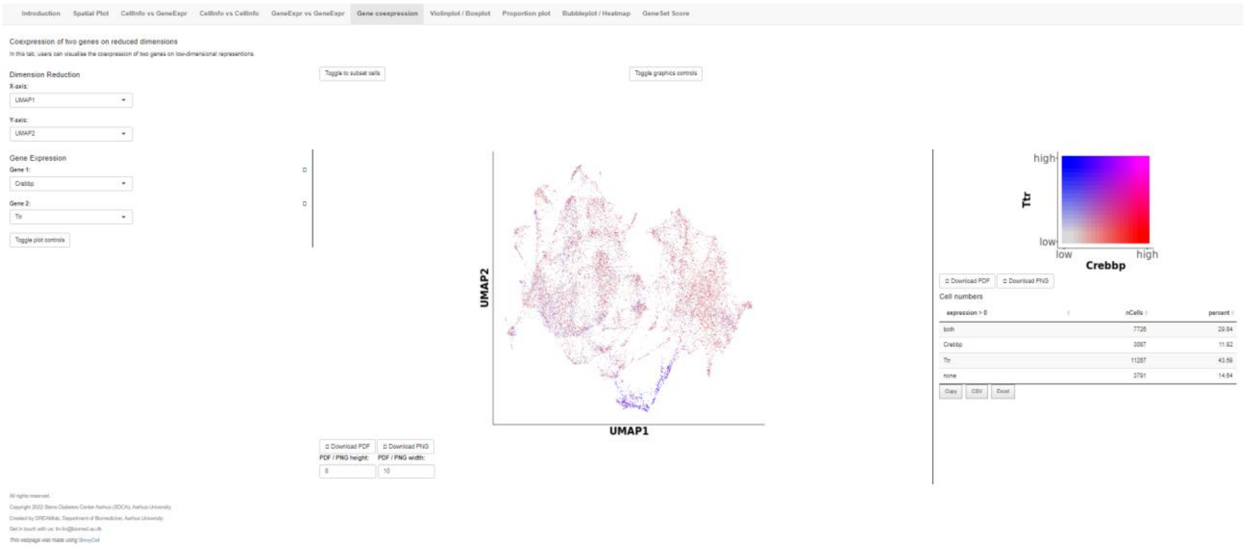

#### 3. Visualizing Gene Expression with Violin and Box Plots

##### <u>3.1. Violin/Box Plot</u>:

- This section provides a visual representation of gene expression distribution using violin or box plots;
- Select the X-axis and Y-axis parameters, and choose the plot type (violin or boxplot);
- Select "Toggle to subset cells" to choose the cell information to subset; Choose between group or annotation to select specific brain regions
- Toggle between showing data points and adjusting graphics controls;
- Download the plots as PDF or PNG files.

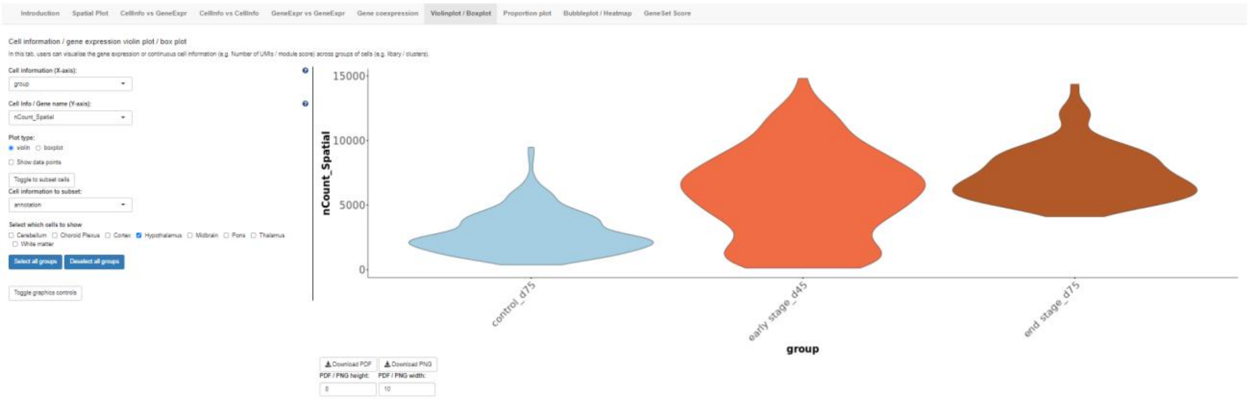

#### 5. Creating Gene Expression Bubbleplots/Heatmaps

##### <u>5.1. Bubbleplot/Heatmap</u>:

- This section allows you to visualize gene expression patterns of multiple genes.
- Enter a list of gene names and select the grouping parameter (e.g., group).
- Choose the plot type (bubbleplot or heatmap).
- Cluster rows (genes) and columns (samples) for better visualization.
- Download the plots as PDF or PNG files.

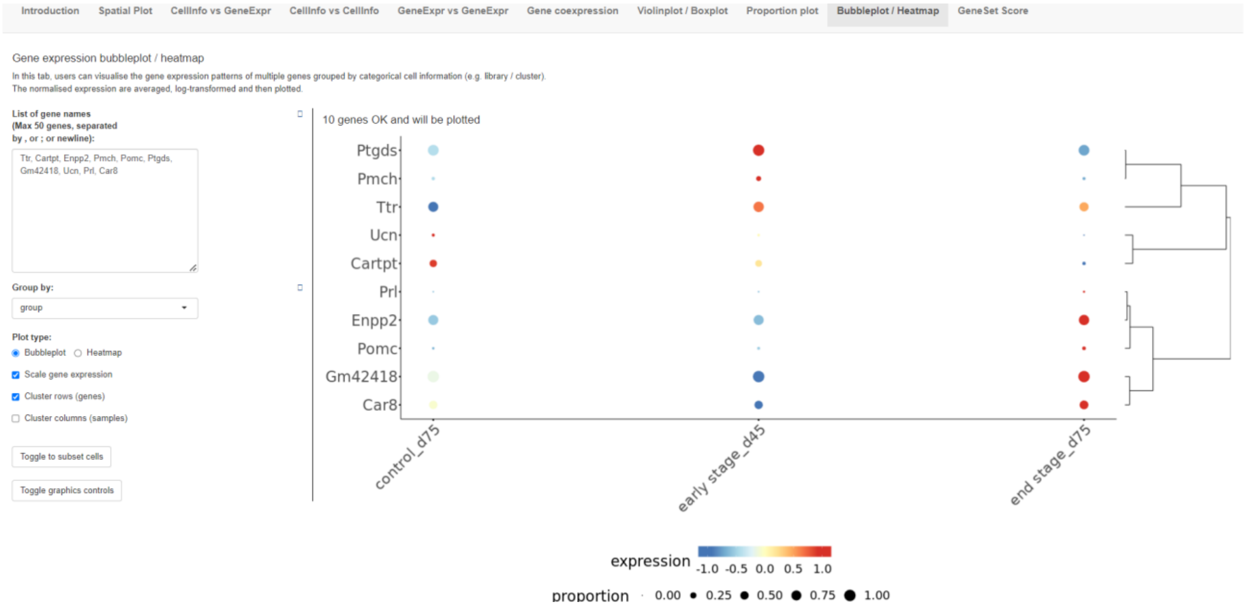

#### 6. How to Use GeneSet Score Tab

The GeneSet Score tab in the Mouse PD spatial database allows users to visualize gene set scores across different brain regions and conditions. This feature helps in understanding the collective behavior of groups of genes involved in specific biological processes or pathways. Here’s a detailed guide on how to use this feature effectively:

##### 6.1. Accessing GeneSet Scores

Click on the "GeneSet Score" tab on the top navigation bar to open the gene set score analysis interface.

##### 6.2. Visualizing GeneSet Scores

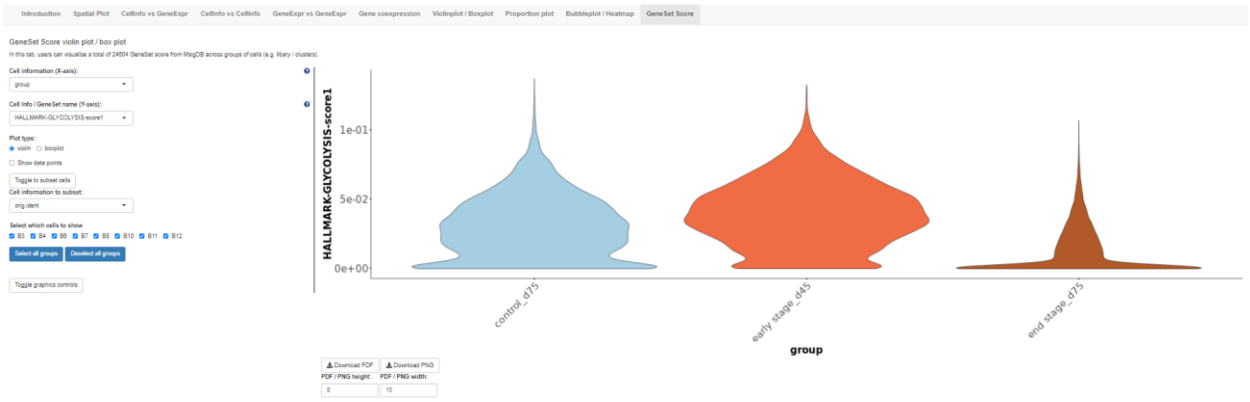


###### 6.2.1. Cell Info/ Gene Set name (Y-axis):

- Use the dropdown menu under "Cell Info/ Gen Set name" to select a gene set/pathway of interest (e.g., "HALLMARK-GLYCOLYSIS-score1");
- Alternatively, you can type keywords related to your gene set of interest in the search box and select from the suggested options;
- Select Plot type for optimal visualization (Violin or boxplot);
- Select “Toggle to subset cells” to choose the cell information to subset; Choose between group or annotation to select specific brain regions;

#### 6.3. Spatial geneset score: Spatial plot (on the spatial plot tab)

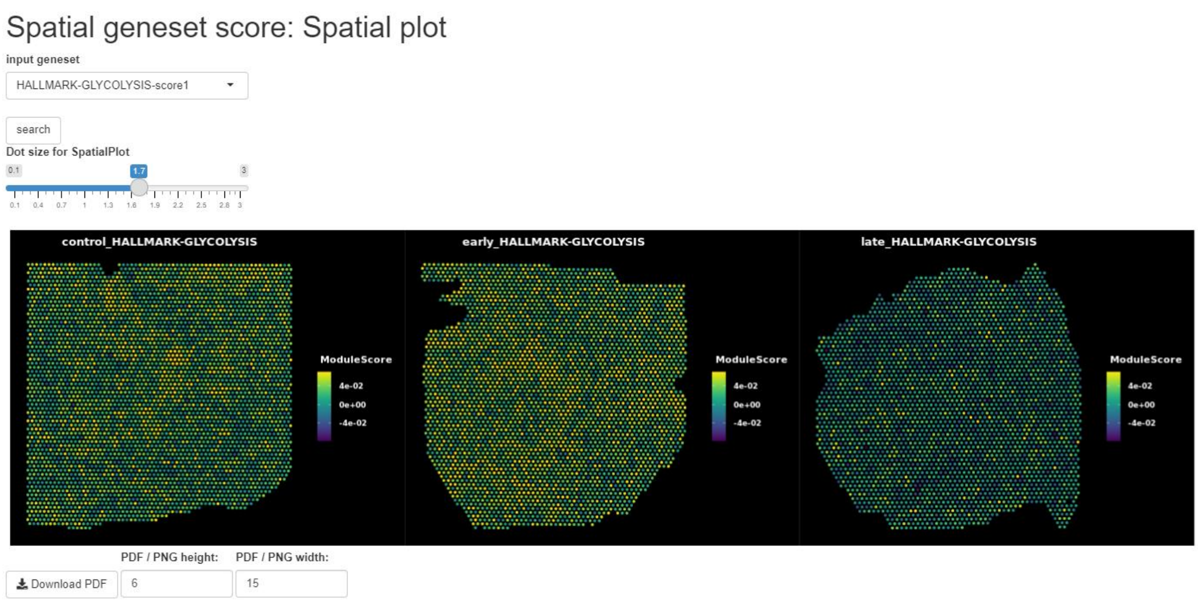


**6.3.1. Input Gene Set**:

- Use the dropdown menu under "input geneset" to select a gene set of interest (e.g., " HALLMARK-GLYCOLYSIS-score1”
- Alternatively, you can type keywords related to your gene set of interest in the search box and select from the suggested options.
- Click the "search" button to retrieve the gene set score data.

**6.3.2. Spatial Plot**:

- The spatial plot provides a visual representation of the gene set scores across different conditions and time points.
- Adjust the dot size for the spatial plot using the slider to change the visualization resolution.
- The example spatial plot displays the gene set score for " HALLMARK-GLYCOLYSIS-score1" across control, early and late PD conditions;
- Download the spatial plot as a PDF or PNG file by setting the desired dimensions and clicking the download buttons.
